# SNP and Haplotype Regional Heritability Mapping (SNHap-RHM): joint mapping of common and rare variation affecting complex traits

**DOI:** 10.1101/2021.08.02.454788

**Authors:** Richard F. Oppong, Thibaud Boutin, Archie Campbell, Andrew M. McIntosh, David Porteous, Caroline Hayward, Chris S. Haley, Pau Navarro, Sara Knott

## Abstract

We describe a genome-wide analytical approach, SNP and Haplotype Regional Heritability Mapping (SNHap-RHM), that provides regional estimates of the heritability across locally defined regions in the genome. This approach utilises relationship matrices that are based on sharing of SNP and haplotype alleles at local haplotype blocks delimited by recombination boundaries in the genome. We implemented the approach on simulated data and show that the haplotype-based regional GRMs capture variation that is complementary to that captured by SNP-based regional GRMs, and thus justifying the fitting of the two GRMs jointly in a single analysis (SNHap-RHM). SNHap-RHM captures regions in the genome contributing to the phenotypic variation that existing genome-wide analysis methods may fail to capture. We further demonstrate that there are real benefits to be gained from this approach by applying it to real data from about 20,000 individuals from the Generation Scotland: Scottish Family Health Study. We analysed height and major depressive disorder (MDD). We identified seven genomic regions that are genome-wide significant for height, and three regions significant at a suggestive threshold (p-value < 1 × 10^−5^) for MDD. These significant regions have genes mapped to within 400kb of them. The genes mapped for height have been reported to be associated with height in humans. Similarly, those mapped for MDD have been reported to be associated with major depressive disorder and other psychiatry phenotypes. The results show that SNHap-RHM presents an exciting new opportunity to analyse complex traits by allowing the joint mapping of novel genomic regions tagged by either SNPs or haplotypes, potentially leading to the recovery of some of the “missing” heritability.

**Author Summary:** In untangling the genetic contribution to observed phenotype differences, situations can arise where causative variants might be tagged by haplotypes and not in linkage disequilibrium with individual SNPs. This scenario is likely for relatively newly arisen and rarer variants. Here, we propose a regional heritability method, SNHap-RHM, that jointly fits haplotype-based and SNP-based genomic relationship matrices (GRMs) to capture genomic regions harbouring rare variants that the SNP-based GRMs might miss. By analysing ^~^20,000 Scottish individuals, we show by simulation that the two GRMs are very specific to the type of variant effects they can capture; – the haplotype-based GRMs specifically target haplotype effects which are mostly missed by SNP-based GRMs and vice versa. Applying the method to height and major depressive disorder led to the uncovering of regions in the genome that harbour genes associated with those traits. These results are uniquely important because first they confirm that effects tagged by haplotypes may be missed by conventional SNP-based methods. Secondly, our method, SNHap-RHM, presents an exciting new opportunity to analyse complex traits by allowing the joint mapping of genomic regions tagged by either SNPs or haplotypes, potentially leading to the recovery of some of the “missing” heritability.

## Introduction

Estimates of the genetic component of complex trait variation using genotyped SNPs led to the conclusion that a proportion of the heritability of complex traits is still unexplained or “missing” (1,2). Full sequence data will contain all the variants that account for all the heritability of complex traits (3). Moreover, some of these true causal variants may be rare (4) and therefore may be in incomplete linkage disequilibrium (LD) with genotyped SNPs (5). Thus, some of the “missing” heritability may be “hidden” in rare variants whose effects are difficult to capture because of lack of statistical power. There is, therefore, some benefit to be gained in terms of improving the heritability estimates and uncovering gene variants involved in the control of traits by fitting genome-wide analytical models that adequately capture the combined effects of rare genetic variants (6,7).

In light of this, we proposed a genome-wide analytical approach that draws its theoretical basis from the genome-based restricted maximum likelihood (GREML) approach (1,2,8–10) which utilises both local and genome-wide relationship matrices to provide regional estimates of the heritability across locally defined regions in the genome (11,12). This regional heritability analysis can capture the combined effect of SNPs in a region, and thus small effect variants may be detectable. However, the analysis only captures effects associated with common SNPs present on genotyping chips.

Haplotypes may provide a better strategy to capture genomic relationships amongst individuals in the presence of causal rare variants. Although rare variants are not in LD with genotyped variants and thus are difficult to capture in conventional GWAS, these rare variants, may be in LD with some haplotypes and thus can be captured using haplotype methods. Compared with genotyped SNPs, capturing haplotype effects may offer an advantage because haplotypes can be functional units (13). Therefore, haplotype effects may reflect the combined effects of closely linked cis-acting causal variants (14) and using haplotypes could provide real benefit over SNPs in recovering some of the “missing” heritability and identifying novel trait-associated variants. Therefore, we extended the SNP-based regional heritability analysis further by incorporating haplotypes in addition to SNPs in the calculation of the regional GRMs used in the analysis (15). This approach includes two regional GRMs and divides the genome into windows based on local haplotype blocks delimited by recombination boundaries.

This paper further explores the properties of both the SNP-based and the haplotype-based regional heritability mapping (SNP-RHM and Hap-RHM respectively). We hypothesise and show by simulation that the Hap-RHM complements existing SNP-RHM analytical approaches by capturing regional effects in the genome that existing SNP-based methods fail to capture. This leads us to propose a mapping strategy that jointly utilises SNP and haplotype GRMs in a single analysis called SNHap-RHM. We then confirm the utility of this approach by applying it to real data obtained from about 20,000 individuals from the Generation Scotland: Scottish Family Health Study (GS: SFHS) (16). We analysed two phenotypes: height and major depressive disorder (MDD). The aim was to uncover novel genetic loci that may affect these traits and improve the estimates of the genetic components of the variation in these traits.

## Results

### Overview of methods

We have shown previously that regional GREML analysis (Regional Heritability Mapping or RHM) using fixed region sizes in the genome is a suitable mapping method for finding local genetic effects (11). The conventional RHM model fits two genomic relationship matrices (GRMs) in the analyses to map genetic loci that affect trait variation: a local GRM (rGRM) calculated using SNPs located in the region and a genome-wide GRM (gwGRM) calculated from SNPs outside the region. We have since extended this conventional regional heritability analysis to incorporate haplotypes in the calculation of the local GRM and have successfully implemented this in a simulation study (15). This study, like our previous (15), utilises a regional heritability model that breaks the genome into naturally defined regions by delimiting them by recombination hotspots. Two types of regional heritability models are then fitted in turn to the phenotypes. One model (SNP-RHM) uses SNPs to estimate local genetic relationships between study individuals, and the other model (Hap-RHM) estimates local genetic relationships amongst individuals using haplotypes.

We first explored the two models in detail using a simulation study in which we simulated 20 replicates of five phenotypes using data from about 20,000 individuals of the GS: SFHS cohort. We then performed a regional heritability analysis that jointly fitted the SNP and the haplotype GRMs in an approach that we termed SNP and Haplotype Regional Heritability Mapping (SNHap-RHM). An overview of SNHap-RHM is shown in Fig 1. We finally applied SNHap-RHM to height and major depressive disorder (MDD) phenotypes of the GS: SFHS.

**Fig 1.**
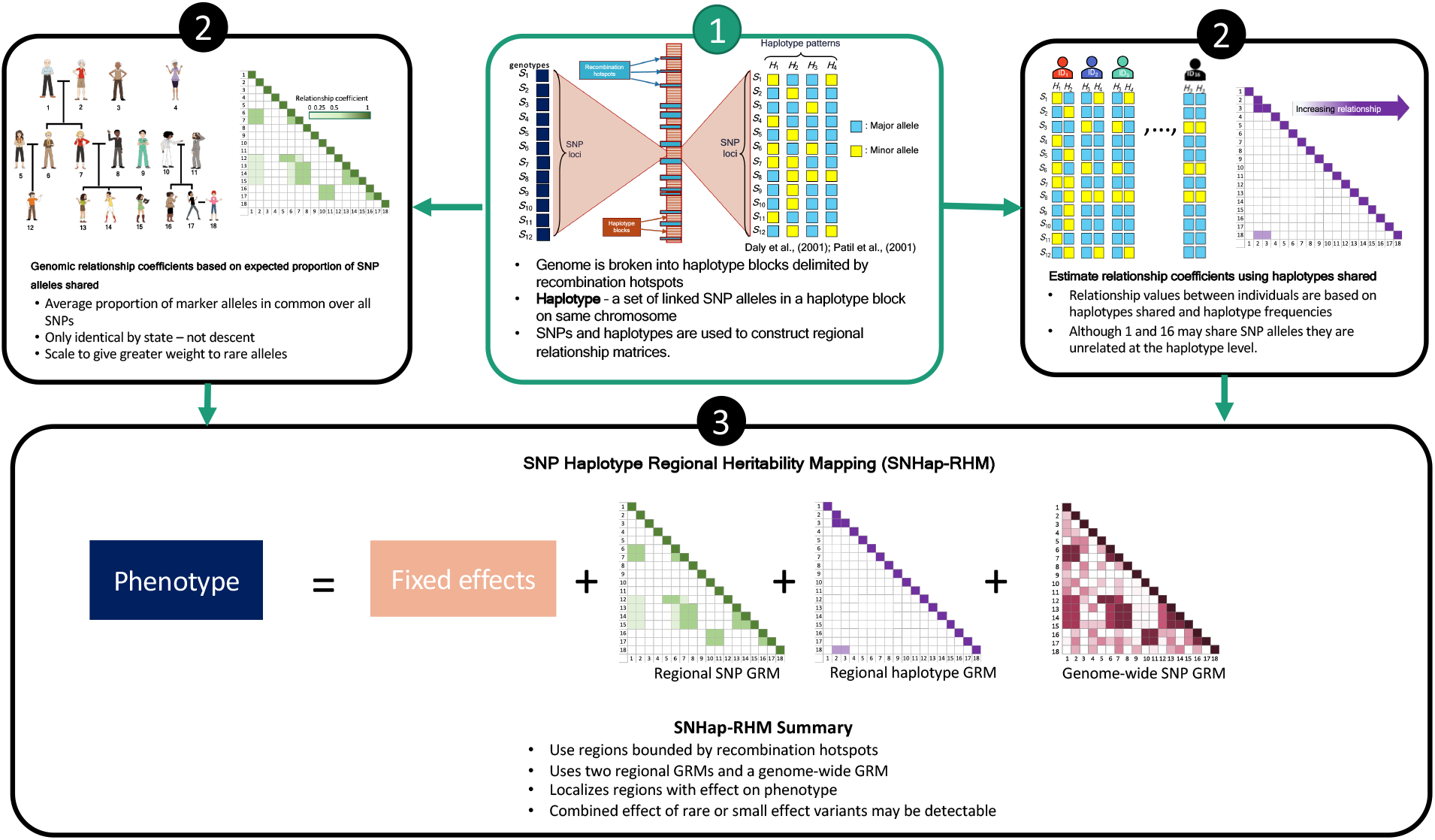
A Schema outlying SNHap-RHM.

Further details of the models, phenotype simulations and GS: SFHS dataset are presented in the materials and methods section of the manuscript.

### Simulation study: SNP-RHM, Hap-RHM and SNHap-RHM

We performed a regional heritability analysis that fits two GRMs (one for the region and one for the rest of the genome) per region across multiple genomic regions delimited by recombination hotspots (where the estimated recombination frequency exceeds ten centiMorgans per Megabase (10cM/Mb)). This recombination threshold resulted in a total of 48,772 regions across the genome. We tested two types of regional heritability models, SNP-RHM and Hap-RHM, on 20 replicates of five simulated phenotypes. In SNP-RHM, the regional matrix is derived from SNP genotypes whereas in Hap-RHM the regional matrix is derived from haplotypes. The phenotypes were simulated to be determined by 20 regional QTL effects and genome-wide polygenic effects. The regional QTL effects of the five phenotypes were simulated using SNPs as causal variants for two of them and haplotypes for the remaining three as described in the methods section.

A likelihood ratio test (LRT) was used to test the null hypothesis, H_0_: that the genetic variance explained by the region is not significant, against the alternative hypothesis, H_1_: that the region accounts for a significant proportion of the phenotypic variance. A large LRT statistic is evidence against the null hypothesis, and therefore means the region explains a significant proportion of the phenotypic variance.

The LRTs averaged over the 20 replicates of the five phenotypes are shown in Fig 2. The figure shows plots of average LRT for the QTL regions and ten adjacent regions (5 to each side). The results show that both models detected the simulated regional effects at the genome-wide significance level (LRT = 23.9) (p-value < 1.02 × 10^−6^, Bonferroni correction for testing 48,772 regions) and can capture true causal loci in traits with different genetic architectures. The LRTs were higher on average for the SNP-based model (SNP-RHM) than the haplotype-based model (Hap-RHM). This could be because for Hap-RHM, the genome-wide GRM which is a SNP-based GRM does not tag any of the background haplotype effects that are outside any one particular region being analysed, and thus the residual variance may be inflated by the other haplotype QTLs which downwardly impact the LRTs.

**Fig 2.**
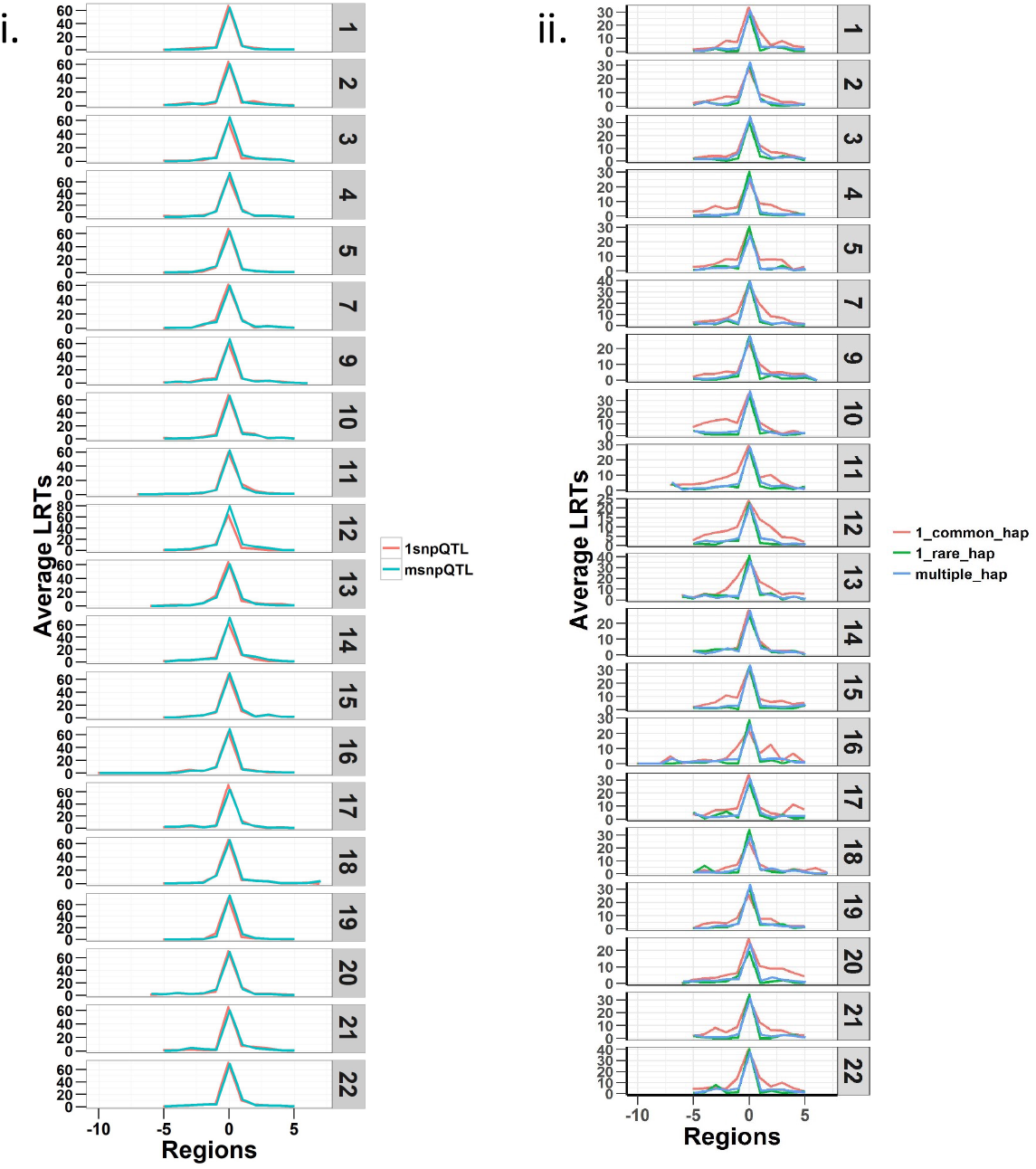
Plots of Likelihood ratio test (LRT) statistics at each QTL locus and 5 regions either side averaged for the 20 simulations of each of the five QTL phenotypes. Plot (i) is SNP QTL phenotypes analysed using the SNP-RHM and plot (ii) is the haplotype QTL phenotypes analysed using the Hap-RHM. Both models can capture the simulated QTL effects for their respective SNP and haplotype phenotypes.

We provide further investigation of the results from the simulation in the supporting information (S1 Text). For both analysis models, we have presented detailed results of the relationships between the LRT statistics, region size, variance estimates and allele frequencies (S3-S10 Figs). We observed that the longer haplotype blocks had many SNPs (and hence many, many haplotypes, up to 14,000 in some blocks), and this impacted the estimation of the simulated regional variance (S8 Fig). We, therefore, performed a hybrid-Hap-RHM analysis that restricted the natural haplotype block sizes to 20 or fewer SNPs per haplotype block. This hybrid-Hap-RHM was to investigate whether the regional variance is well captured by Hap-RHM when shorter haplotypes are used. The hybrid-Hap-RHM underestimated the regional variance for larger regions but did not offer any discernible improvement in the LRT statistics (S9 Fig). The relationship between region size and estimated variance was different between the Hap-RHM and hybrid-Hap-RHM, while we observed a similar relationship between LRTs and the region size.

Both SNP-RHM and Hap-RHM fail to capture the simulated regional effects when the simulated phenotype has a genetic architecture that does not match the analysis model, i.e., SNP or haplotype (Fig 3 and S1 Fig). These figures show the results for the situation where the SNP QTL phenotypes were analysed with the haplotype-based model (Hap-RHM) and the haplotype QTL phenotypes were analysed with the SNP-based model (SNP-RHM). Both models fail to detect the simulated effects in such situations, therefore, showing that the models complement each other since they capture effects due to different types of genetic variants (i.e., tagged by SNPs or haplotypes).

**Fig 3.**
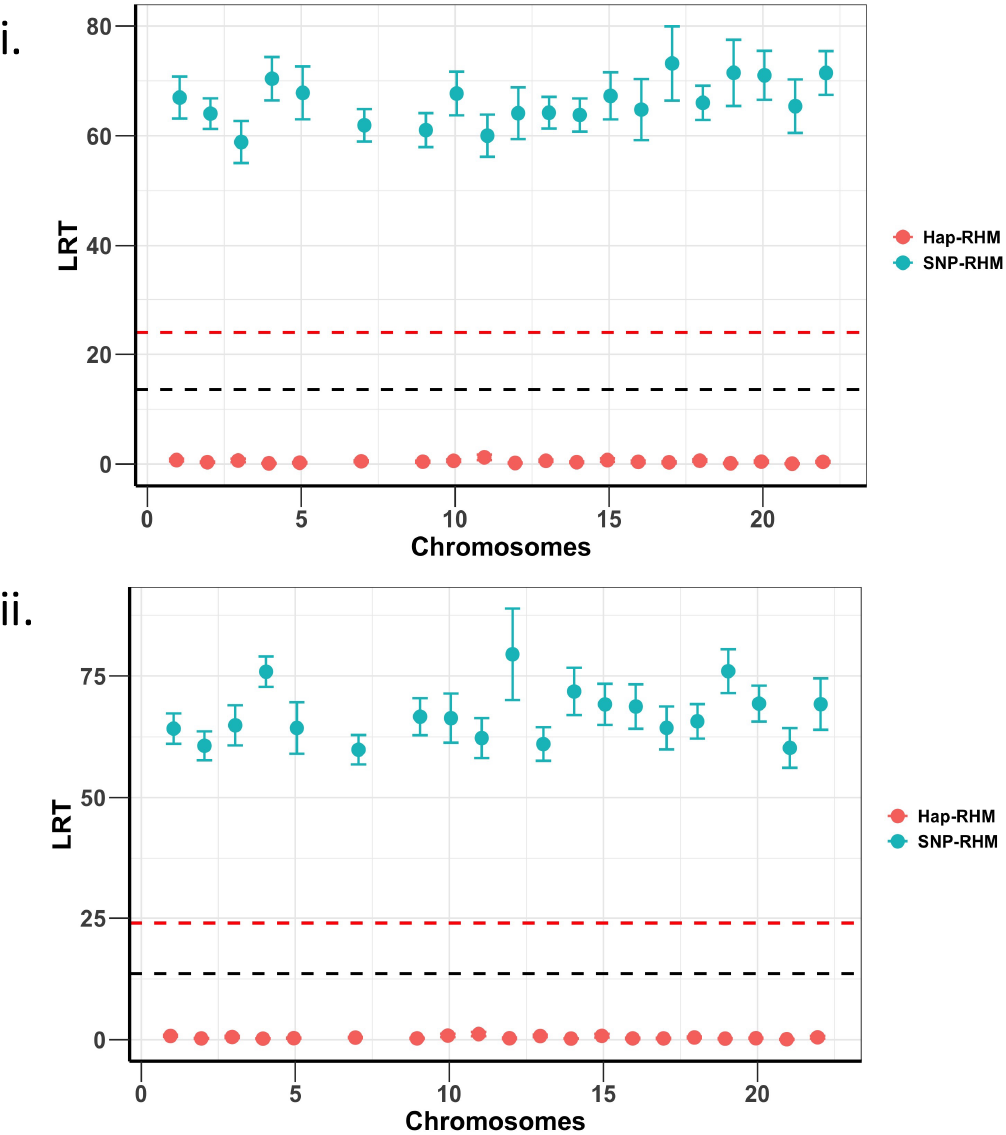
Plots of average LRT statistics over replicates of QTL loci across the chromosomes for the 20 simulations of each of the two SNP QTL phenotypes. The red dashed lines are genome-wide significance threshold (for 48,772 regions) and the black dashed lines are Bonferroni significance threshold (for 220 regions). The upper plot (i) is the 1-SNP QTL phenotype, and the lower plot (ii) is the multiple SNP QTL phenotype. The two phenotypes are analysed using both the SNP based model (SNP-RHM) (blue points) and the Haplotype based model (Hap-RHM) (red points). The Hap-RHM fails to capture the simulated effects for the SNP QTLs.

To confirm that two models are complementary and independent of each other, we implemented SNHap-RHM that fits the regional SNP and haplotype GRMs jointly, on a replicate of each of the five simulated phenotypes. The significance of regional effects was tested with an LRT with two degrees of freedom. The results are shown in Fig 4 and confirm that the two models are complementary since even when we fitted jointly the two regional matrices (SNP and Haplotype-based), we can still capture the simulated effects.

**Fig 4.**
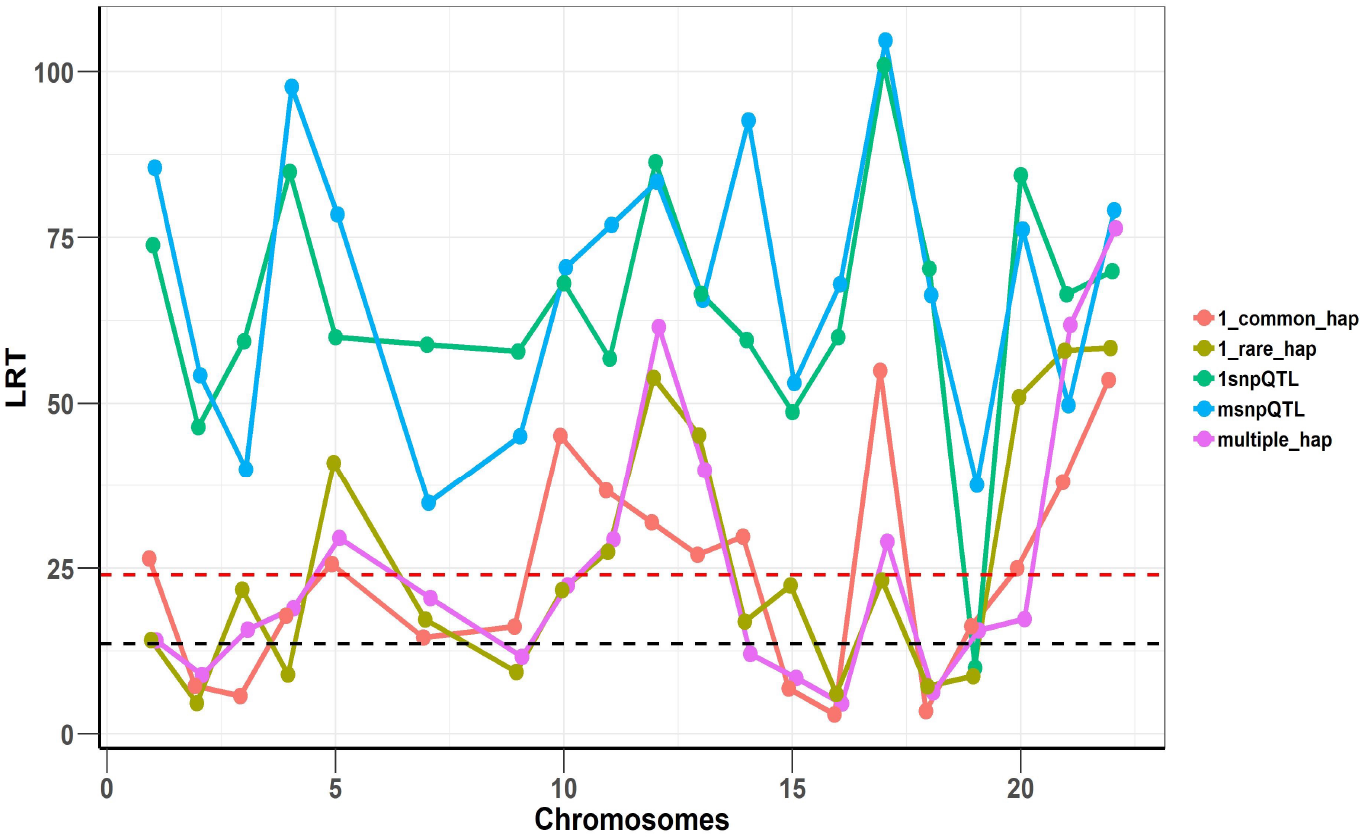
Joint analysis of the SNP and haplotype phenotypes using SNHap-RHM. The plot is an analysis of one replicate of each of the simulated phenotypes. The LRT statistics are plotted over QTL loci across the chromosomes. The red dashed lines are genome-wide significance threshold (for 48,772 regions) and the black dashed lines are Bonferroni significance threshold (for 220 regions).

### SNHap-RHM analysis of height and MDD in GS: SFHS

The heritability estimates for height and MDD in the GS: SFHS dataset, calculated using the whole-genome GRM, were 81.4% (0.92) and 13.8% (1.35) respectively. There were no overlaps between regions identified as significant (tested with an LRT with one degree of freedom) by the haplotype and SNP-based models for either of the two traits (S2 Fig). This reaffirms our hypothesis tested by simulation that the Hap-RHM is complementary to SNP-RHM in mapping associated genomic loci.

The regional heritability results for height and MDD are presented as plots of minus-Log10 of the LRT p-values (Figs 5 and 6). The plots for the SNHap-RHM, SNP-RHM and Hap-RHM analyses are shown.

**Fig 5.**
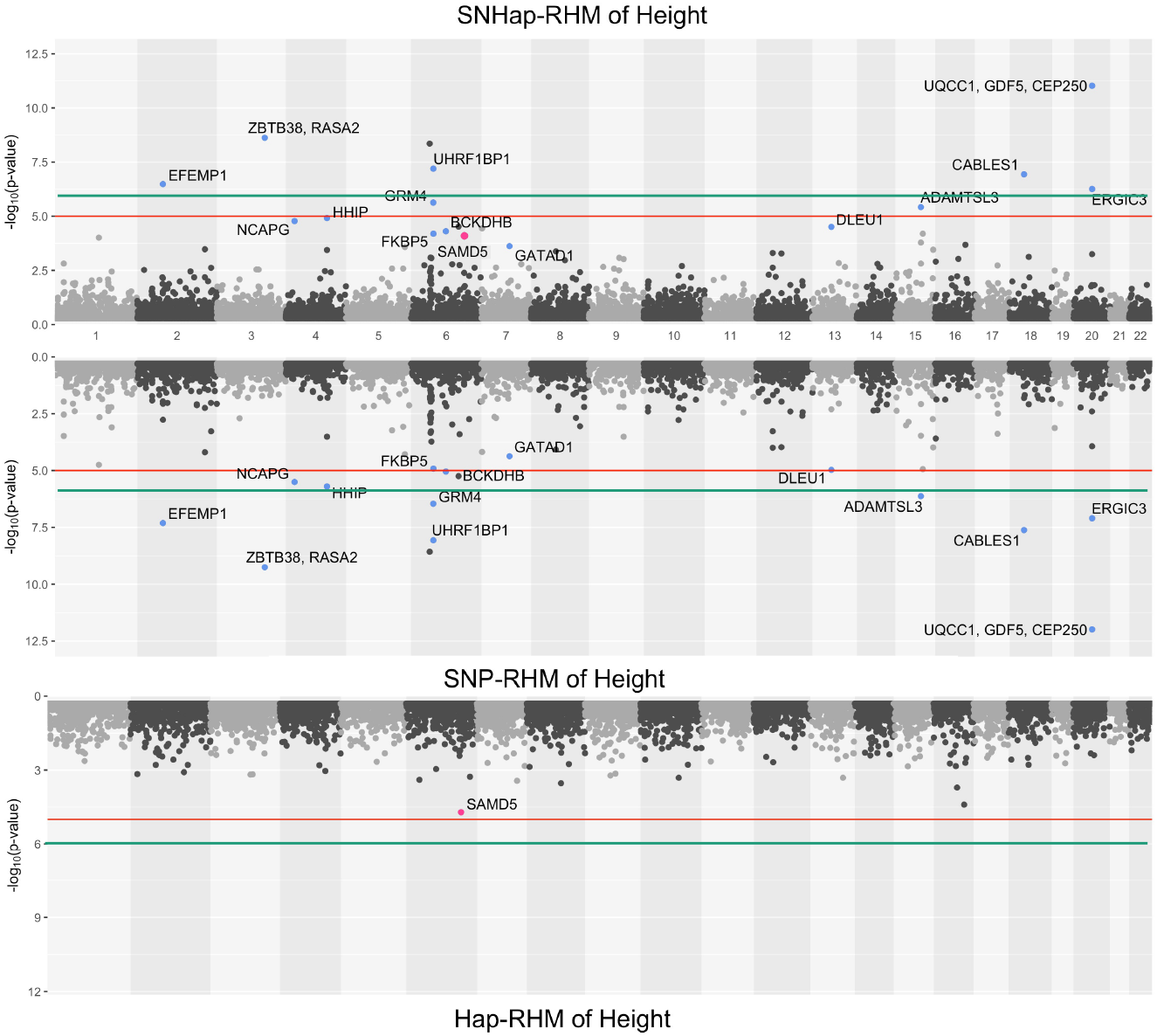
The genome-wide evidence of haplotype block association for height. Analysis done with SNHap-RHM, SNP-RHM and Hap-RHM. The points are plots of -log10 of the p-values of regions tested with the LRT for the regional GREML analyses. The green lines are the Bonferroni-corrected genome-wide significance threshold and the red lines are the suggestive significance threshold calculated to be p-value < 1 × 10^−5^. The top association hits at p-value < 5 × 10^−5^ with genes located within the region are highlighted in blue for SNP-RHM and red for the Hap-RHM.

**Fig 6.**
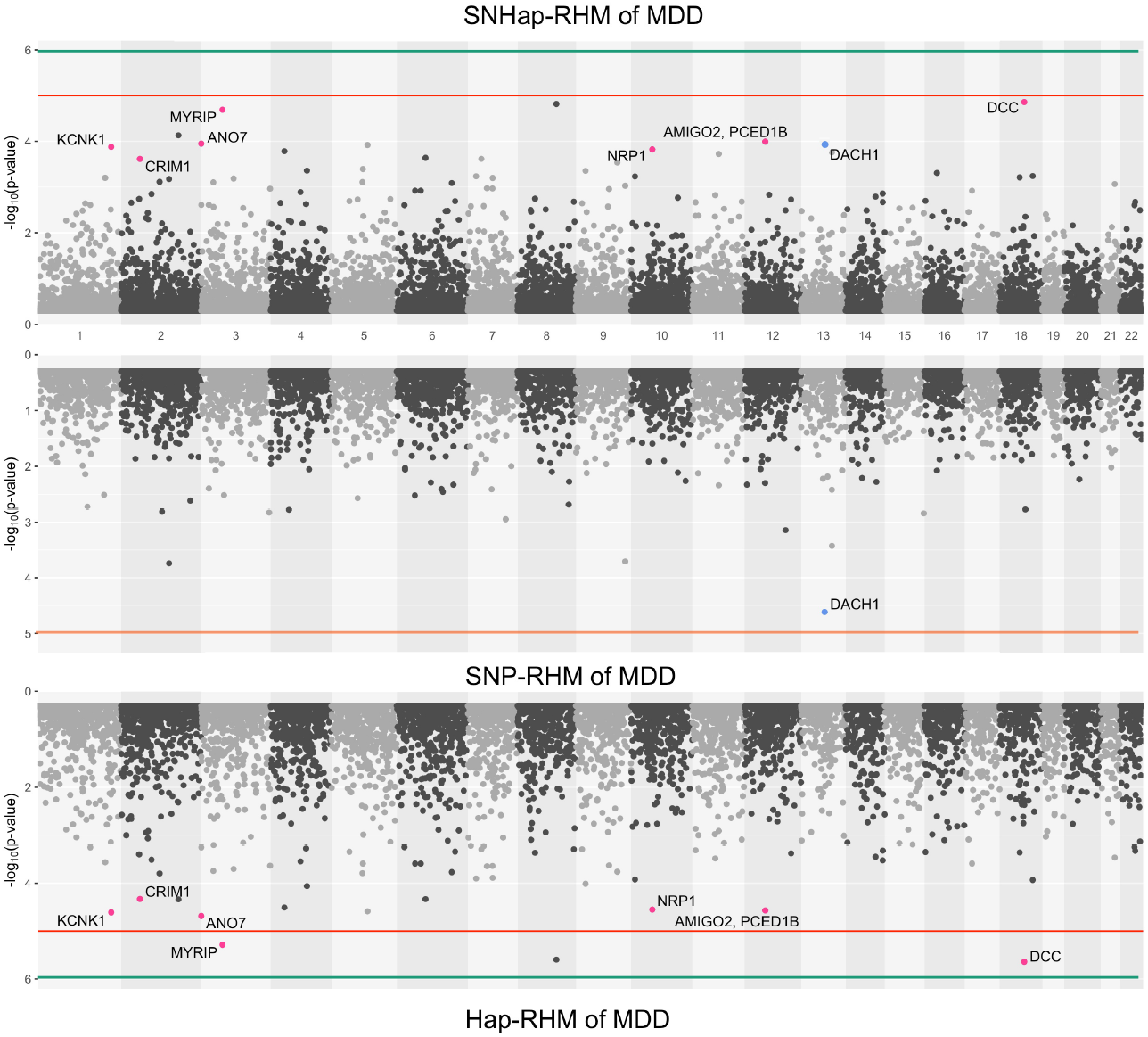
The genome-wide evidence of haplotype block association for Major Depressive Disorder. Analysis done with SNHap-RHM, SNP-RHM and Hap-RHM. The points are plots of -log10 of the p-values of regions tested with the LRT for the regional GREML analyses. The green lines are the Bonferroni-corrected genome-wide significance threshold and the red lines are the suggestive significance threshold calculated to be p-value < 1 × 10^−5^. The top association hits at p-value < 5 × 10^−5^ with genes located within the region are highlighted in blue for SNP-RHM and red for the Hap-RHM.

The results for height show that nine regions passed the Bonferroni-corrected genome-wide significance threshold in the analysis using SNP-RHM. No region was genome-wide significant for height when analysed with Hap-RHM. Furthermore, seven of the nine associated regions still come up as genome-wide significant when SNPs and haplotypes in those regions are analysed jointly using SNHap-RHM. There are GWAS reported genes that lie in or are within 400kb of these regions (S1 Table).

For MDD, no region passed the Bonferroni-corrected genome-wide significance threshold for the analysis done with the SNP-based and haplotype-based regional heritability models (Fig 6). Three regions passed the suggestive significance threshold at p-value < 1 × 10^−5^ for Hap-RHM analysis of MDD. A further nine regions were significant at p-value < 5 × 10^−5^ for the haplotype-based analysis, and one region for the SNP-based analysis (S2 Table). Figure 6 shows that when the two local GRMs are fitted jointly using SNHap-RHM, the genomic regions associated with MDD can still be mapped. The associated regions mapped by the haplotype-based model for MDD contain genes reported by GWAS to be associated with several psychiatry phenotypes (Fig 6 and S2 Table). The most strongly associated region was within 400kb of the *DCC* gene. This gene is part of the NETRIN1 pathway, which has been reported to be associated with major depressive disorder in two GWAS samples (GS: SFHS and Psychiatric Genomics Consortium) (17). Zeng *et al*. (17) used a SNP-RHM guided by pathway analysis (to first uncover pathway association and then localise *DCC* within the pathway) to show the *DCC* association with major depressive disorder. The second most strongly associated region was on chromosome 8, and this region had no gene mapped to it.

A linear mixed effects model was used to test for association of the SNPs within the suggestive significant region identified by the haplotype-based model on chromosome 3 for MDD. The model tested for association of SNPs by fitting their allelic dosages individually in a regression model and fitting a GRM to account for relatedness of individuals. The region on chromosome 3 was chosen in this example because there is a psychiatric phenotype associated gene, *MYRIP* (18), mapped to it, unlike the *DCC* region which has the gene outside the region. The results are shown in Table 1. Five SNPs within this region are nominally significant at p-value < 0.05. Four out of these five SNPs confer about 2% increased risk of the disease each. These four SNPs lie within the *MYRIP* gene sequence. The *MYRIP* gene is expressed in the brain (19). A SNP (rs9985399) in this gene is reported to be associated with brain processing speed in the Lothian birth cohort (18). Brain processing speed is an important cognitive function that is compromised in psychiatric illness such as schizophrenia and depression, and old age. Also, a SNP (rs6599077) in the *MYRIP* gene region is associated with sleep duration (20). Sleep durations outside the normal range (both short sleep and long sleep) is significantly associated with increased risk of depression (21–24). The *MYRIP* gene is also reported to have a role in insulin secretion (25) and low insulin levels have been linked to depression (26–28).

**Table 1.** SNP-based association test of MDD in the *MYRIP* gene region.

| SNP information |  |  |  | Major Depressive Disorder association |  |  |  |
| --- | --- | --- | --- | --- | --- | --- | --- |
| SNP ID | Chr | Pos | MAF | OR | Log (OR) | SE (logOR) | p |
| rs9842160 | 3 | 39844703 | 0.14 | 0.97 | -0.030 | 0.013 | 0.02 |
| rs9858242 | 3 | 39847606 | 0.19 | 1.02 | 0.025 | 0.011 | 0.03 |
| rs1599902 | 3 | 39954674 | 0.41 | 1.02 | 0.019 | 0.009 | 0.04 |
| rs7618607 | 3 | 39947936 | 0.41 | 1.02 | 0.019 | 0.009 | 0.04 |
| rs9860916 | 3 | 39944942 | 0.41 | 1.02 | 0.019 | 0.009 | 0.04 |

The columns are the SNP ID, chromosome, genome position of SNP, minor allele frequency, odds ratio, log of odds ratio, standard error of log odds ratio and association p-value.

#### Comparison with published GWAS SNPs

For both traits, the SNPs in the regions that were significant at p-value < 5 × 10^−5^ were compared to SNPs reported in the GWAS catalogue (29) to be significant for the two traits. The GWAS catalogue was accessed on the 15^th^ of January 2021. The results are presented in Table 2. The SNP-based and haplotype-based models identified 1,380 and 45 SNPs respectively for height, and 78 and 495 SNPs respectively for MDD taking all SNPs within haplotype blocks significant at p-value < 5 × 10^−5^. Out of the 1,380 SNPs identified for height by the SNP-based model, 57 SNPs spanning 20 haplotype regions were in common with published GWAS results for height.

**Table 2.** Comparison of SNPs within significant regions identified by both models and published GWAS results for height and MDD.

| Trait | Number of SNPS |  |  | Number of overlapping SNPS |  |  |
| --- | --- | --- | --- | --- | --- | --- |
|  | SNP-RHM | Hap-RHM | pubGWAS | SNP-RHM & Hap-RHM | SNP-RHM & pubGWAS | Hap-RHM & pubGWAS |
| Height | 1380 | 45 | 4960 | 0 | 57 | 0 |
| MDD | 78 | 495 | 1815 | 0 | 0 | 0 |

The columns are the name of trait, number of SNPS in regions identified by SNP-RHM and HAP-RHM with $p\text{-value} < 5 \times 10^{-5}$ and SNPS in published GWAS (pubGWAS) for the traits, and the number of SNPS overlapping between the three.

## Discussion

We have proposed and implemented a genome-wide analytical method that analyses genomic regions using a regional heritability model (11). We have since extended this method to include haplotypes by fitting a regional haplotype-based GRM (Hap-RHM) and redefined genomic regions in our analysis to be delimited by recombination hotspots generated using HapMap Phase II (15,30). In this study, we build on our previous regional heritability methods by exploring the properties of the SNP and haplotype-based regional heritability mapping models by simulation and demonstrate that the two variance components fitted are largely independent of each other (S2 Fig). The novelty in this study is that we show that the two regional matrices fitted in SNP-RHM and Hap-RHM capture two different kinds of effects in terms of genetic architecture, and thus the two variance components can be fitted jointly (by fitting the SNP and haplotype regional matrices together) in a joint marker regional heritability mapping procedure that we call SNHap-RHM.

We hypothesised that the Hap-RHM would complement the SNP-RHM. We investigated this hypothesis in a simulation study in which we simulated 20 replicates each of two types of SNP QTL phenotypes and three types of haplotype QTL phenotypes. The results show that the two heritability models can capture the effects of causal variants within genomic loci associated with the phenotype analysed. The results also show that the two models are specific about the type of causal effect they can capture, therefore, providing support for the hypothesis that haplotype-based regional heritability models will complement SNP-based regional heritability models. We provide further support for this hypothesis by fitting the two GRMs jointly and showing (using an LRT with two degrees of freedom) that we can still capture the simulated effects and real effects from real data.

We applied SNHap-RHM to height and MDD phenotypes from the Generation Scotland: Scottish Family Health Study. Again, we draw comparisons between the effects captured by the SNP-RHM and the Hap-RHM. The SNP-RHM identified more Bonferroni-corrected genome-wide (GW) significant regions (p-value < 1.02 × 10^−6^) for height compared to MDD. Fifty-seven of the SNPs identified for height by the SNP-RHM have been reported by other studies to be associated with height. These SNPs spanned 20 genomic regions in the GS: SFHS cohort. Height is a highly polygenic trait with many common genetic variants accounting for most of the additive genetic variation (31). These common genetic variants may be in LD with genotyped SNPs on SNP chips (these chips are disproportionately enriched for common SNPs). Therefore, the SNP-based regional heritability model is better suited for capturing SNP loci in height compared to MDD.

MDD is a very heterogeneous phenotype, and thus every MDD case could have a set of genetic and non-genetic risk factors exclusive to them (32). These unique genetic risk factors will mean that a lot of the genetic variants driving the disease will be rare at the population level. Three genomic regions were identified for MDD by the haplotype-based regional heritability model at the suggestive level, p-value < 1 × 10^−5^. The Hap-RHM works well for MDD because MDD is believed to be driven by rare genetic variants, and the model can capture rare genetic variants. The haplotype model can capture rare variants because of the LD between rare variants (both typed and untyped) and the flanking variants that aggregate to form the haplotypes within the genomic regions. There were no overlaps between regions identified by the Hap-RHM and SNP-RHM for each trait, which again supports the hypothesis that the two models complement each other in mapping associated loci.

In both traits, the top significant regions we mapped at p-value < 5 × 10^−5^ had genes mapped to those regions or within 400kb of those regions. For height, these genes have been reported to be associated with height in humans (33–39). For MDD, these genes have been reported to be associated with major depressive disorder and other psychiatry phenotypes (17,18,40–43). In one of such regions for MDD, five SNPs within the region are individually significantly associated with MDD at the nominal level (p-value < 0.05). Four of these SNPs lie within the gene sequence of *MYRIP*, and they each confer 2% disease risk. A conventional GWAS analysis would have missed these nominally associated SNPs because they will not reach the suggestive significance threshold, let alone genome-wide (GW) significance. However, analysing these SNPs within the region as haplotypes allowed us to detect the combined effect of these SNPs in the region at a suggestive-significance level even with our relatively small sample size compared to recent genome-wide association studies of MDD: 322,580 (44) and 480,359 (43).

The current study’s primary strength is that we show the ability of SNHap-RHM to incorporate SNP and haplotype information jointly to map genomic regions that affect complex traits. This gives SNHap-RHM a uniquely useful role to play in the future of complex traits analysis. The plummeting costs of whole-genome resequencing (45) have shifted research focus in GWA studies towards sequence data analysis (46). Although whole-genome sequence data analysis allows incorporating all the genetic variants that drive the phenotypic variation, there may still be some variants whose individual effects may be too small to be picked up in a conventional GWA analysis. However, regionally analysing sequence information can help overcome this because multiple small-effect variants in a region can add up to a substantial regional effect that can be captured by a regional SNP GRM or tagged by a haplotype GRM. Moreover, by defining haplotype blocks using recombination hotspots, whole-genome information can be summarised naturally without setting an arbitrary number of SNPs, and that facilitates integration and comparison across studies. More so, regional heritability analysis of sequence data would be an efficient way to deal with the burden of multiple testing, which has long been a problem of conventional GWAS.

One limitation of the current study is the computation burden of the analyses, which necessitates the pre-correction of the phenotypes with the whole-genome GRM before performing SNHap-RHM. This was a leave-one-chromosome-out step involving 22 separate GREML analyses, each fitting a whole-genome GRM that excluded SNPs from one chromosome (47). For our sample of about 20,000 individuals, the precorrection step reduced the computation time needed to perform GREML analysis at each region by approximately 33% (15 minutes) and used about 20% (16 gigabytes) less memory. Although this was done to speed up the analysis, the precorrection step was used as an approximation to account for the background polygenic effects of genetic markers outside each region; this would have been about 48,772 separate GREMLs to account for each region. Also, due to the two degrees of freedom test applied in SNHap-RHM, we observed a slight drop in the significance of the associated regions in both height and MDD when SNHap-RHM was applied to those traits. One option would be to use a less stringent test for SNHap-RHM, effectively testing regions assuming only one degree of freedom so that if only one of the variance components significantly contributed to the phenotypic variance the region would be identified for subsequent formal testing of the individual variance components.

Finally, although this study thoroughly evaluates the robustness of SNP and Haplotype RHM using simulation and demonstrates the utility of SNHap-RHM in real phenotype analysis, seeking replication in other cohorts will improve our understanding and, more importantly, demonstrate that the analysis is portable across studies and genotyping platforms.

## Conclusion

We have implemented a regional heritability analysis and undertaken analyses of regions in the genome delimited by recombination boundaries and shown by simulation that haplotype-based GRMs can capture genetic variance that may be missed by conventional SNP-based GRMs. We then applied this method in the analysis of real phenotype data from GS: SFHS. Again, we show that the haplotype-based regional heritability model uncovers associations in regions of the genome that explain genetic variance missed by the SNP-based heritability model. In light of this, we further showed that regional effects can still be captured when the two regional GRMs (SNP and haplotype-based) are fitted jointly: an analytical procedure we termed SNHap-RHM. This SNHap-RHM presents an exciting new opportunity to analyse complex traits by allowing the joint mapping of novel genomic regions tagged by either SNPs or haplotypes, potentially leading to the recovery of some of the “missing” heritability.

## Materials and Methods

### Ethics Statement

Ethical approval for the GS: SFHS study was obtained from the Tayside Committee on Medical Research Ethics (on behalf of the National Health Service).

### The general statistical setting of a regional heritability analysis

Consider a vector ***y*** of phenotype values with length *n*, the linear mixed-effects model for fitting the effects of genomic region *i* and background polygenic markers is given as:

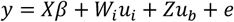

where ***y*** is a vector of phenotypes, ***X*** is a design matrix of fixed effects, and ***β*** is a vector of fixed effects, ***W_i_*** is a design matrix relating phenotype measures to genetic markers in region *i* and ***u_i_*** is a vector of random genetic effects due to region *i* assumed to be multivariate normal, 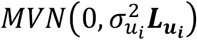. ***L_u_i__*** is a relationship matrix calculated using markers (SNPs or haplotypes) in region *i*: calculated in the subsequent sections as ***G*** for the SNP and ***H*** for the haplotype-based models. ***Z*** is a design matrix for background polygenic effects of markers outside the region *i* and ***u_b_*** is a vector of random polygenic effect of genetic markers excluded from region *i*, assumed to be multivariate normal, 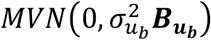. ***B_u_b__*** is a relationship matrix calculated using the markers outside the region *i*: calculated in the subsequent section in the same way as ***G***. And ***e*** is a vector of residual effects assumed to be multivariate normal, 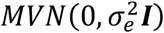. ***I*** is an identity matrix.

Under the model, the vector of phenotypes ***y*** is assumed to be normally distributed, *N*(***Xβ, V***) where the variance is

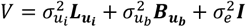

#### SNP-RHM: SNP-based regional heritability model

A SNP-based regional heritability analysis was first reported by Nagamine *et al*. (11). The regional heritability analysis approach we employ here differs from the analysis done by Nagamine *et al*. (11) in the way the regions are defined. That analysis defined local regions by breaking the genome into smaller user-defined windows of *p* SNPs, which overlapped by *q* SNPs. Here, however, we define regions based on recombination boundaries in the genome.

The regional heritability model fits two genetic relationship matrices (GRMs): one local GRM for the region and a whole-genome GRM for the remaining SNPs in the genome that are outside the region. The GRMs are genomic relatedness matrices calculated as the weighted proportion of the local or genome-wide autosomal SNPs shared identity by state (IBS) between pairs of individuals. The SNP IBS matrices are calculated as follows, following the second scaling factor proposed by VanRaden (48)

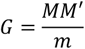

where *m* is the total number of *r* local or *b* background autosomal SNPs, and ***M*** is a matrix of genotype codes for the sampled individuals centred by loci means and normalised by the standard deviation of each locus. ***M*** is calculated as follows for individual *i* at locus *j*

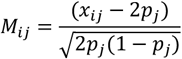

where *x_ij_*, is the genotype code at locus *j* for individual *i* and takes the values 0, 1 and 2 for AA, Aa and aa genotypes respectively, *p_j_* is the frequency of allele ‘a’ at locus *j*. The SNP-based relationship for individuals *i* and *k* is therefore calculated as follows

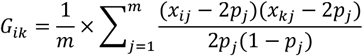

#### Hap-RHM: Haplotype-based regional heritability model

The haplotype-based regional heritability model follows theoretically from the SNP-based analysis and utilises haplotypes instead of SNPs as the genetic markers for the regional analysis. The analysis fits two GRMs, a haplotype-based regional GRM and a SNP-based background genome-wide GRM. The haplotype-based GRM is similar to the SNP-based GRM defined in the previous section. For a locally defined region (haplotype block) containing *h* haplotype variants, the haplotype-based kinship for individuals *i* and *k* is calculated as follows

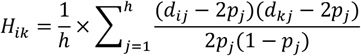

where *d_ij_*, is the diplotype code (coded as the number of copies of haplotype *j*) for individual *i* and takes the values 0, 1 and 2 for the *h_t_h_t_, h_t_h_j_, h_j_h_j_* diplotypes respectively where haplotype *t* is any haplotype other than haplotype *j*, i.e. *t* ≠ *j, p_j_* is the haplotype frequency for haplotype *j*.

### Phenotype simulations

Five phenotypes were simulated using available genotypic information of 20,032 individuals from the Generation Scotland: Scottish Family Health Study (16). A total of 593,932 genotyped SNPs were used, and missing genotypes were filled in by imputation. A total of 555,091 SNPs remained after a QC that removed SNPs of MAF < 0.01 and SNPs that were out of Hardy-Weinberg equilibrium at p-value < 0.000001.

The five phenotypes were simulated to have a total variance of 1. This total is composed of 0.6 environmental (residual) variance and genetic variance of 0.4. The genetic variance was partitioned into two components, a polygenic variance of 0.3 and a total QTL variance of 0.1 (20 QTLs, each explaining a variance of 0.005). A common polygenic variance was simulated for all five phenotypes from 20,000 markers randomly selected across the genome. The polygenic variance was simulated to be normally distributed with zero mean and variance of 0.3.

For each phenotype, 20 regions (haplotype blocks) were randomly selected, one on each autosome (except chromosomes 6 and 8 because of the unusually high LD in the MHC regions on chromosome 6 and a large inversion on chromosome 8 (49)), to simulate quantitative trait loci (QTL). This gave a total of 20 QTLs for each phenotype. The regions were delimited by natural boundaries: recombination hotspots where the estimated recombination frequency exceeds ten centiMorgans per Megabase (10cM/Mb) with the estimated recombination frequency between boundaries being less than ten centiMorgans per Megabase (10cM/Mb) based on the Genome Reference Consortium Human Build 37 (50). This recombination threshold resulted in a total of 48,772 regions across the genome. The number and type of marker used to simulate the QTL are what defined the five phenotypes. The five phenotypes are, a 1-SNP QTL within the haplotype block, a multiple-SNP (5 SNPs) QTL within the haplotype block, two types of 1-haplotype QTL within the haplotype block (taking either a common or a rare haplotype as causal) and multiple (5) haplotype QTL within the haplotype block. Details of these phenotypes are described below.

For the haplotype QTL phenotypes, a haplotype block is treated as a single genetic locus having multiple alleles. Each haplotype variant within a block is considered as an allele of that locus. Each study individual will carry two alleles, or have a diplotype, for each locus or haplotype block. The genotype data used to simulate the phenotypes were phased using SHAPEIT2 (51) to produce the haplotypes for study individuals. The multiple haplotype QTL phenotypes were simulated by randomly sampling two rare haplotypes and three common haplotypes within each haplotype block to give five haplotypes per block. The two types of 1-haplotype QTL phenotypes were simulated by randomly sampling a rare haplotype per haplotype block for one type and for the other type a common haplotype was randomly sampled within each haplotype block. S10 Fig gives an indication of the frequencies for the rare (0.00002 to 0.036) and common haplotype (0.008 to 0.906) randomly sampled to simulate the phenotypes. There is a slight overlap between the frequencies for rare and common haplotypes because the regions had already been randomly selected before proceeding to randomly select rare and common haplotypes in those regions. Which means what is rare in one region may be common in another.

The individual marker contribution to the polygenic effect and the QTL effects were calculated as follows

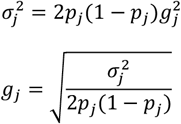

where 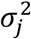 is the contribution of a marker to the QTL or polygenic variance, *g_j_* is the effect of a SNP *j* or haplotype *j* randomly sampled to have polygenic or QTL effect, *p_j_*, is the frequency of haplotype *j* or the effect allele of the SNP *j*. For the single marker QTL phenotypes, each QTL explained a variance of 0.005. For the multiple marker QTL phenotypes, each causal variant explained the same variance, with the effects scaled to account for LD in the region so each QTL locus explained a variance of 0.005. For the multiple haplotype QTL effects, the haplotype effects were scaled relative to the inverse of their frequency to give a total variance explained by the region of 0.005.

Common environmental effects were randomly sampled for the five phenotypes from a normal distribution 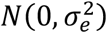 where 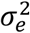 is 0.6. This, together with a genetic variance of 0.4, gave a total variance of 1 for each phenotype. The final simulated phenotype for an individual *i* was then calculated as follows

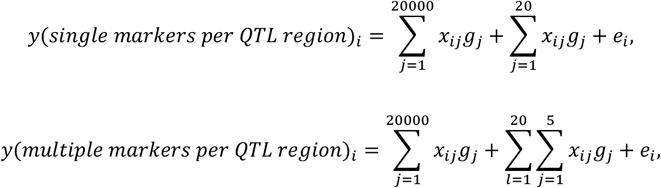

where *x_ij_*, is the number of copies of the effect allele of SNP *j* for individual *i* (for haplotypes, this is defined as *d_ij_*; the number of copies of haplotype *j* for individual *i*) and *g_j_* is the effect of haplotype *j* or SNP *j*. Twenty replicates were analysed for each of the five phenotypes with a different set of QTL markers sampled for each replicate.

#### Analysis of simulated data

The five simulated phenotypes were analysed using the two models, the SNP-based regional heritability model (SNP-RHM for the SNP QTL phenotypes) and the haplotype-based regional heritability model (Hap-RHM for the haplotype QTL phenotypes). To test the analytical models’ specificity, we applied Hap-RHM to SNP QTL phenotypes and SNP-RHM to the haplotype QTL phenotypes. We also performed a Hap-RHM analysis in which the units of analysis in the haplotype blocks were restricted to regions of 20 or fewer SNPs per haplotype block. This was because we observed that longer haplotype blocks had many SNPs (and hence many, many haplotypes, up to 14,000 in some blocks), and this impacted the estimation of the simulated regional effect. The hybrid Hap-RHM, therefore, investigates whether the regional effect is well captured by the haplotype-based model when shorter haplotypes are used.

We estimated the regional genetic variance and polygenic variance using restricted maximum likelihood (REML). For each simulated phenotype, we analysed 220 regions in total to map the 20 simulated QTLs. This involved analysing the region containing the QTL and ten adjacent regions (five in either direction). In this way, we limit the analysis to the regions in the genome with simulated effects, thereby reducing computation time considerably. Also, by analysing neighbouring regions, we are able to explore the precision of estimates of the location of regional effects. We assessed the significance of a region using the Likelihood Ratio Test (LRT). The genome-wide significance threshold was calculated to be LRT = 23.9 (p-value < 1.02 × 10^−6^) using a Bonferroni correction for testing 48,772 regions.

Also, we selected one replicate for each simulated phenotype and performed a regional heritability analysis that jointly fitted the SNP and the haplotype GRM in an approach that we termed SNP and Haplotype Regional Heritability Mapping (SNHap-RHM).

### GS: SFHS Data

#### Genotyping, quality control and phasing of Generation Scotland: Scottish Family Health Study dataset

The data from the Generation Scotland: Scottish Family Health Study (GS: SFHS) comprised 23,960 participants recruited from Scotland (16,52). The DNA from about 20,032 of the participants had been genotyped using the Illumina HumanOmniExpressExome8v1-2_A chip (^~^700K genome-wide SNP chip) (16). GRCh37 was used throughout.

Quality control excluded SNPs and individuals with a call rate less than 98%, SNPs with minor allele frequency (MAF) less than 1%and SNPs that were out of Hardy-Weinberg equilibrium (p-value < 0.000001). A total of 555,091 autosomal SNPs passed quality control for downstream analysis. Phasing of the GS: SFHS data was done using SHAPEIT2 (51). Best guess haplotypes were used. Haplotype blocks were defined using recombination hotspots with a recombination rate of 10cM/Mb inferred from the Reference Consortium Human Build 37 (50). Haplotypes variants within blocks were determined using the phased data.

#### Phenotype definition

MDD status for GS: SFHS participants was assigned following an initial mental health screening questionnaire with the questions: “Have you ever seen anybody for emotional or psychiatric problems?” or “Was there ever a time when you, or someone else, thought you should see someone because of the way you were feeling or acting?” Participants who answered yes to one or both of the screening questions were further interviewed by the Structured Clinical Interview for DSM-IV (SCID) (53). A total of 18,725 participants (2,603 MDD cases and 16,122 controls) were retained for analysis for MDD. A total of 19,944 participants from the GS: SFHS were analysed for height.

#### SNHap-RHM of MDD and Height

SNHap-RHM fits jointly, the two types of regional GRMs, SNP-based and haplotype-based, in the analysis of phenotypes (Fig 1). We pre-corrected the phenotypes with the whole-genome GRM before performing SNHap-RHM to speed up the GREML analysis of each block. This pre-correction has previously been shown to speed the regional heritability analysis by Shirali *et al*. (15). This is a leave-one-chromosome-out step (47), which involved 22 separate GREML analyses each fitting a whole-genome GRM that excluded SNPs from one chromosome. The residuals from the pre-correction step were then used in the SNHap-RHM analysis. The models adjusted for sex, age, age^2^, and the first 20 principal components calculated from the study participants’ genomic relationship matrix (calculated using 555,091 autosomal SNPs).

The significance of a region was tested with a likelihood ratio test (LRT) with two degrees of freedom which compared a model with three variance components fitted (the two regional variances together with the residual variance) against a model with only the residual variance component fitted. The individual regional variance components in all regions were subsequently tested with an LRT with one degree of freedom which compared a model with three variance components fitted against a model with two variance components fitted (one regional variance component dropped from the model).

The p-values obtained from the LRTs were used to generate genome-wide association plots for each phenotype (equivalent to GWAS Manhattan plots). The genome-wide significance threshold was calculated to be LRT = 23.9 (p-value < 1.02 × 10^−6^) using a Bonferroni correction for testing 48,772 regions. The suggestive significance threshold of a region was set at an LRT = 19.5 (p-value < 1 × 10^−5^).

## Supporting information

Supplementary results

## Supporting information

S1 Text. Investigating the SNP-RHM and Hap-RHM with simulated phenotypes.

S1 Fig. Plots of average LRT statistics over replicates of QTL loci across the chromosomes for the 20 simulations of each of the three haplotype QTL phenotypes.

S2 Fig. The two analysis models (SNP-RHM and Hap-RHM) are independent of each other in the analysis of height, and Major depressive disorder

S3 Fig. Plots of LRT statistics against QTL region size for the 20 simulations (not averaged) of each of the two SNP QTL phenotypes

S4 Fig. Plots of LRT statistic against QTL region size for the 20 simulations of each of the three haplotype QTL phenotypes.

S5 Fig. Plots of LRT statistic against estimated regional variance for the 20 simulations of the single SNP QTL phenotype.

S6 Fig. Plots of LRT statistic against estimated regional variance for the 20 simulations of each of the three haplotype QTL phenotypes.

S7 Fig. Plots of region size against estimated regional variance for the 20 simulations of the two SNP QTL phenotype.

S8 Fig. Plots of region size against estimated regional variance for the 20 simulations of the three haplotype QTL phenotype.

S9 Fig. Plots for the 1-rare haplotype QTL phenotype analysed using Hap-RHM (red points) and a hybrid variant of the Hap-RHM (blue points).

S10 Fig. Plots of LRT statistic against QTL marker frequencies.

S1 Table. Top genomic regions identified by SNP/ haplotype-based model for Height.

S2 Table. Top genomic regions identified by SNP/ haplotype-based model for MDD.

## Acknowledgements

We are grateful to all the families who took part, the general practitioners, and the Scottish School of Primary Care for their help in recruiting them, and the whole Generation Scotland team, which includes interviewers, computer and laboratory technicians, clerical workers, research scientists, volunteers, managers, receptionists, healthcare assistants and nurses. We also acknowledge Eilidh Fummey for coming up with the name for the joint mapping method.

## Author Contributions

Conceived and designed the experiments: RFO PN CSH SK. Provided data: TB AC AMM DP CH. Performed the experiments: RFO. Analysed the data: RFO. Wrote the paper: RFO PN CSH SK.

