## Supplementary results for "SNP and Haplotype Regional Heritability Mapping (SNHap-RHM): joint mapping of common and rare variation affecting complex traits"

### Supporting Information

#### S1 Text. Investigating the SNP-RHM and Hap-RHM with simulated phenotypes

##### LRT vs Region size

The LRT statistics decayed with increasing region size (when measuring size as number of markers in the region) for the two models even when the analysis models (SNP-RHM or Hap-RHM) matched the genetic architecture of the simulated phenotypes (S3 and S4 Figs). The rate of decay was more pronounced for Hap-RHM, in which the LRT had a non-linear relationship with the region size (S4 Fig).

##### LRT vs Variance estimates

For the SNP QTL phenotypes, the variance estimated by the SNP-RHM in regions where we simulated effects was significantly different from zero (except two regions in both phenotypes) (S5 Fig). The plots show strongly positive curvilinear relationship between the LRTs and the estimated variance. As the LRT increases, the variance approaches and then crosses the simulated variance. This is to be expected and is true for real data as well (e.g., the winner's curse is a similar phenomenon).

On the other hand, the relationship between the LRT and regional variance estimates is less clear for the haplotype QTL phenotypes (S6 Fig). The plots, again, show a non-linear relationship, and once the LRT passes the Bonferroni-corrected threshold for genome-wide significance (red horizontal dashed lines), the variance estimates approached the simulated value of 0.005 (blue vertical line).

#### Region size vs Variance estimates

S7 Fig shows a curvilinear relationship, albeit not significant, between number of markers in the region and the estimated regional variance for the SNP QTL phenotypes. However, the relationship is very strong for the haplotype QTL phenotypes (S8 Fig). The estimated regional variances are inflated when a region has more than 5,000 different haplotype alleles. These regions harbour lots of rare haplotypes, which means the haplotypes chosen as QTLs in the simulation are more likely to be rare. Therefore, the inflation may be an artefact of estimation procedure, which indirectly assumes that rare variants with lower MAF will have larger allele effects and thus by design overestimates their effect. This happens because the GREML model normalises the marker genotypes and assumes that the effect size per normalised genotype follows a normal distribution.

#### LRT vs Allele frequencies

We further investigated whether the allele frequencies of the QTLs influence the LRT statistics and the estimated regional variances. We show in S10 Fig that there is no relationship between allele frequencies of the SNP QTLs and LRTs. There is, however, a linear and curvilinear relationship between LRTs and the haplotype frequencies of the rare and common haplotype QTLs, respectively. The haplotypes simulated to have an effect in regions with an overestimated regional variance are marked with different colours on these plots. These haplotypes are shown to have rare haplotype frequencies and relatively low LRTs (S10 Fig). These results also show that the QTLs for the 1-SNP QTL phenotypes and one common haplotype QTL phenotypes are evenly distributed across the MAF spectrum, whereas the QTLs for the one rare haplotype QTL phenotypes have a MAF distribution that is skewed towards zero.

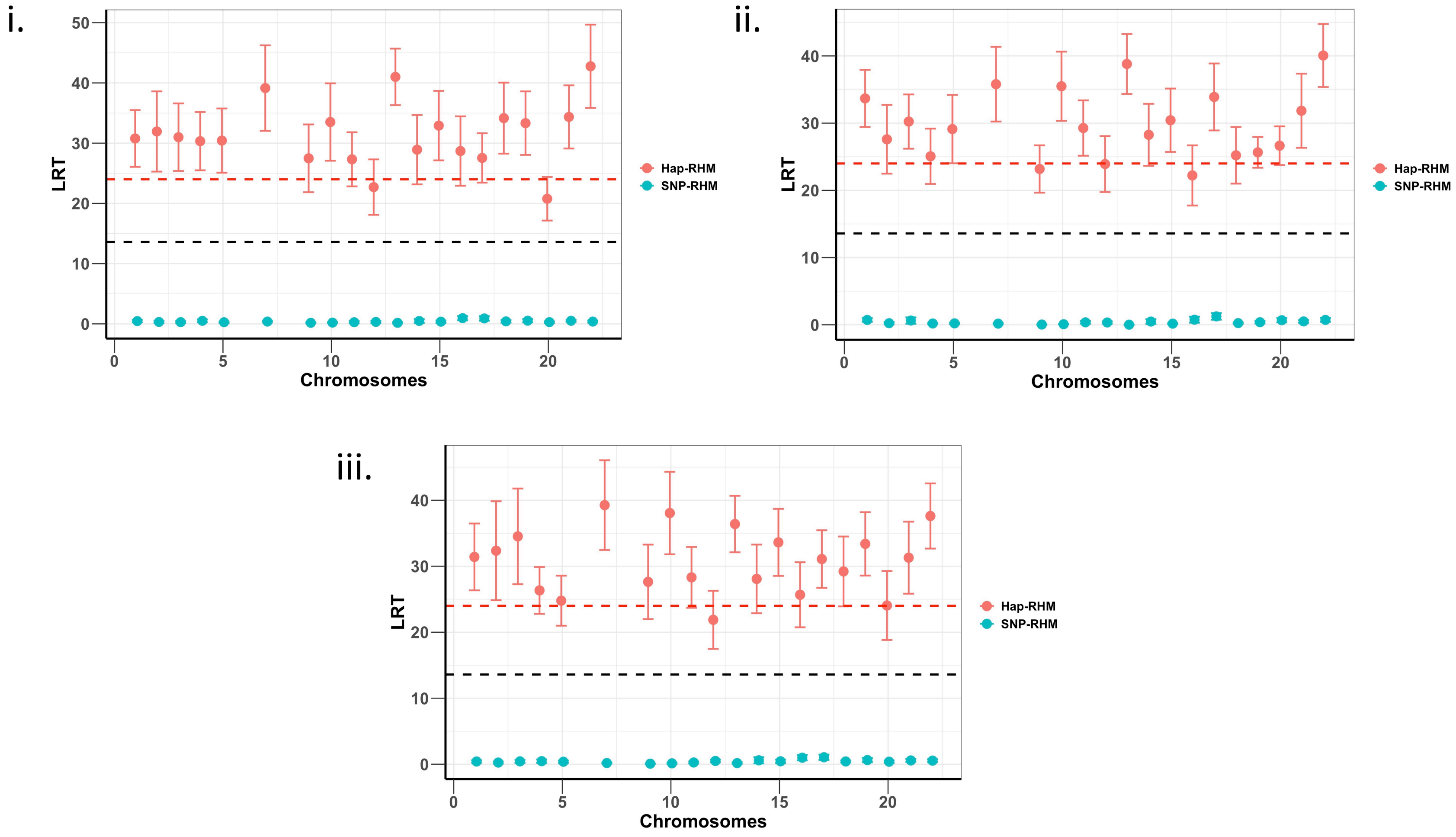


**S1 Fig. Plots of average LRT statistics over replicates of QTL loci across the chromosomes for the 20 simulations of each of the three haplotype QTL phenotypes.** The red dashed lines are genome-wide significance threshold (for 48,772 regions) and the black dashed lines are Bonferroni significance threshold (for 220 regions). The plot (i) is the 1-rare haplotype QTL phenotype, the plot (ii) is the 1-common haplotype QTL phenotype, and the plot (iii) is the multiple haplotype QTL phenotype. The three phenotypes are analysed using both the SNP based model (SNP-RHM) (blue points) and the Haplotype based model (Hap-RHM) (red points)The SNP-RHM fails to capture the simulated effects for the haplotype QTLs.

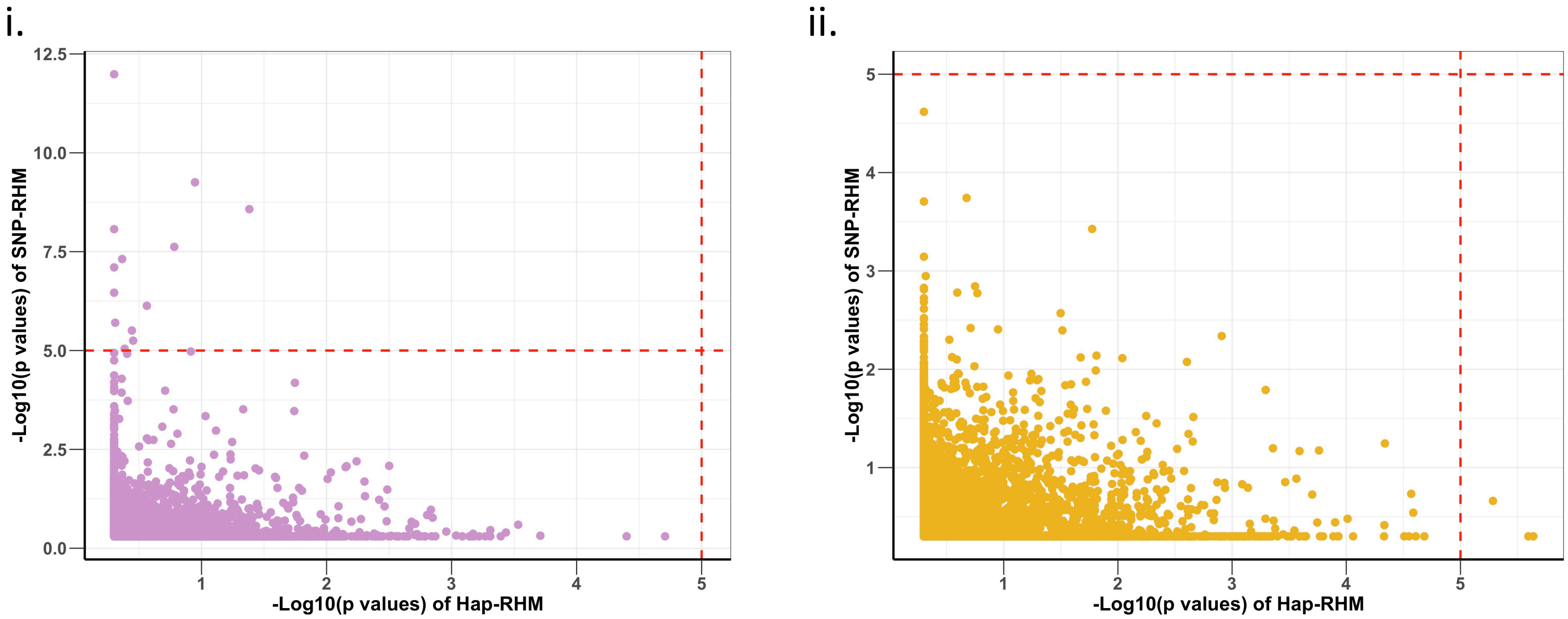


**S2 Fig. The two analysis models (SNP-RHM and Hap-RHM) are independent of each other in the analysis of i. height, and ii. Major depressive disorder.** The minus log10 of p values of association of regions for the SNP-RHM are plotted against that of the Hap-RHM. The red vertical and horizontal dashed lines are the suggestive significance threshold at p-value $< 1 \times 10^{-5}$. The plots show that different significant regions are identified by each model for both phenotypes. The correlation of the regional associations between the two models are 0.11 for height and 0.16 for MDD.

i.

ii.

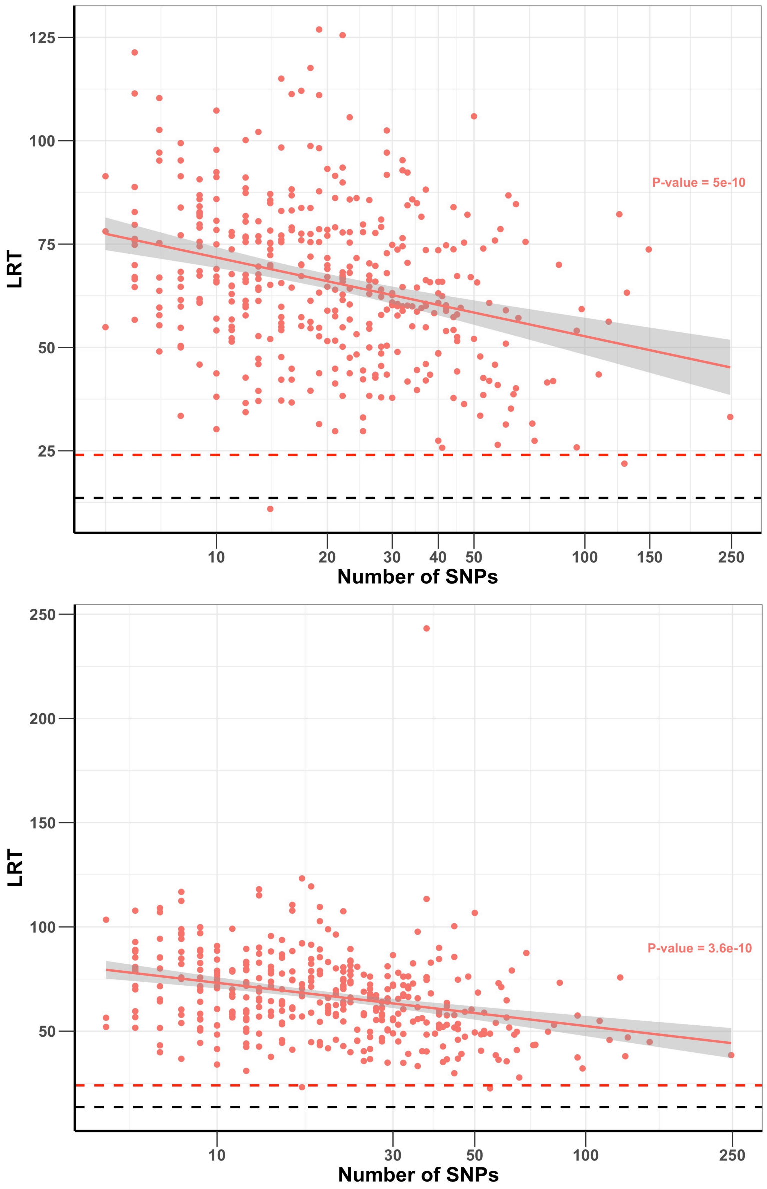


**S3 Fig. Plots of LRT statistics against QTL region size for the 20 simulations (not averaged) of each of the two SNP QTL phenotypes**. The red dashed lines are genome-wide significance threshold (for 48,772 regions) and the black dashed lines are Bonferroni significance threshold (for 220 regions). Plot (i) is the 1-SNP QTL phenotype, and the lower plot (ii) is the multiple SNP QTL phenotype. In both phenotypes the LRT statistic reduced with increasing region size.

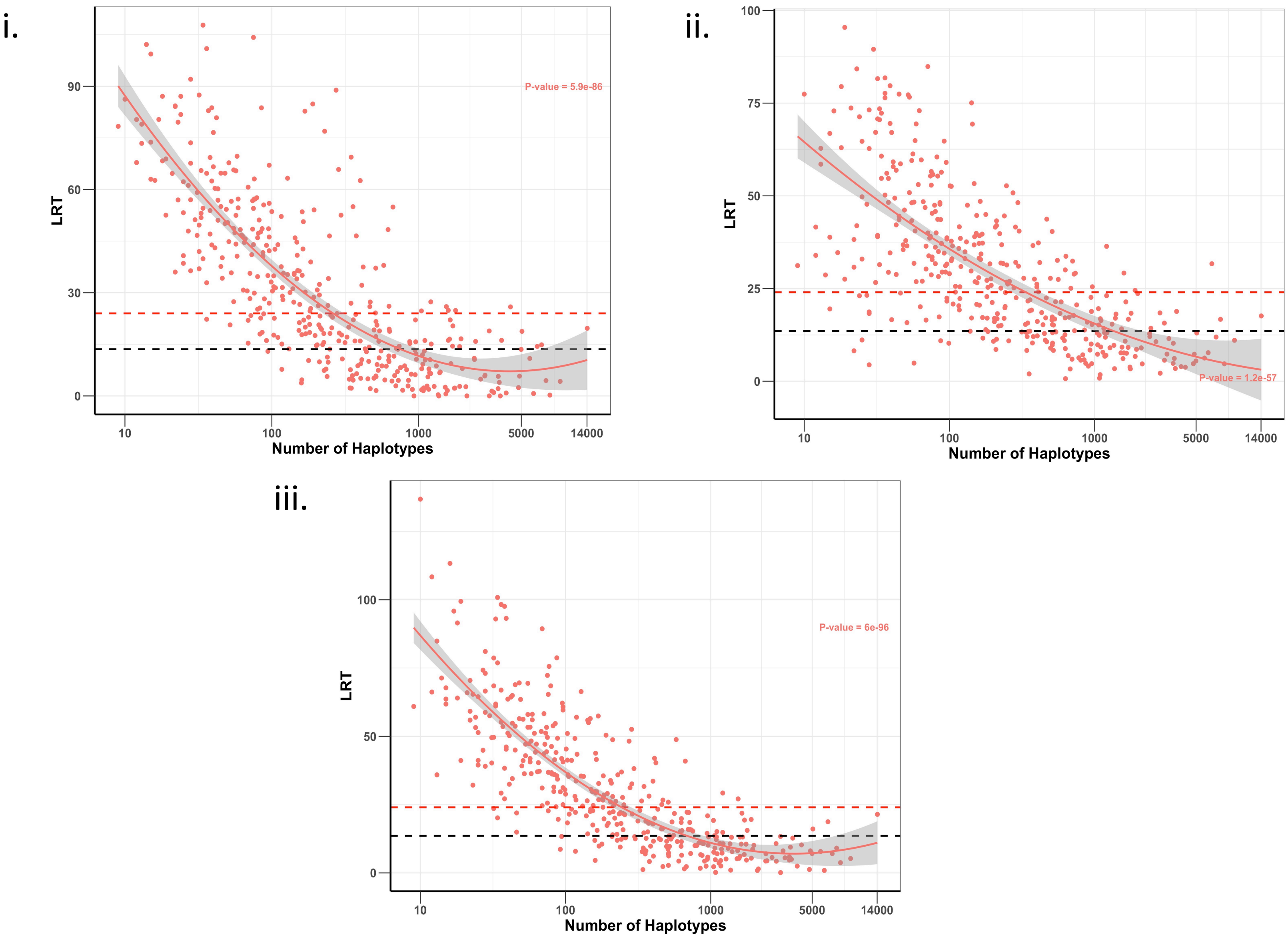


**S4 Fig. Plots of LRT statistic against QTL region size for the 20 simulations of each of the three haplotype QTL phenotypes.** The red dashed lines are genome-wide significance threshold (for 48,772 regions) and the black dashed lines are Bonferroni significance threshold (for 220 regions). The plot (i) is the 1-rare haplotype QTL phenotype, the plot (ii) is the 1-common haplotype QTL phenotype, and the plot (iii) is the multiple haplotype QTL phenotype. For all the three phenotypes the LRT statistic reduced with increasing region size.

i.

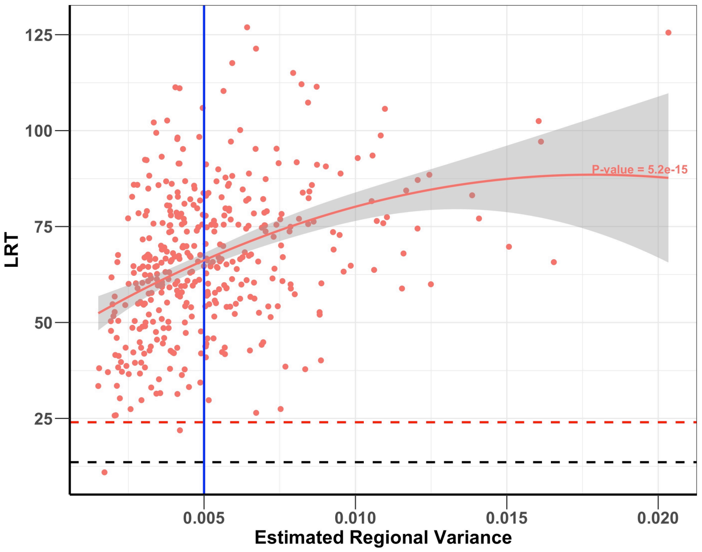


ii.

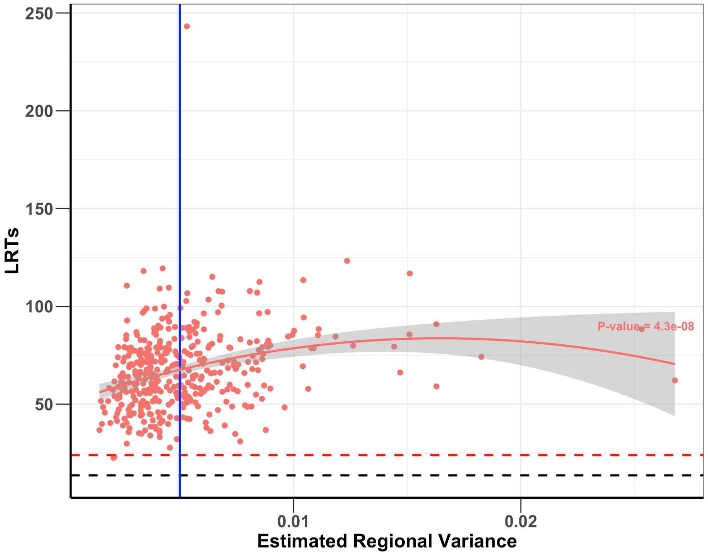


**S5 Fig. Plots of LRT statistic against estimated regional variance for the 20 simulations of the single SNP QTL phenotype.** The red dashed lines are genome-wide significance threshold (for 48,772 regions), the black dashed lines are Bonferroni significance threshold (for 220 regions), and the blue vertical line is the simulated regional variance of 0.005. The plot (i) is the 1 – SNP QTL phenotype, and the lower plot (ii) is the multiple SNP QTL phenotype. The estimated regional variance clustered closely around the simulated value for most of the regions. The variances of a few regions were overestimated in both phenotypes.

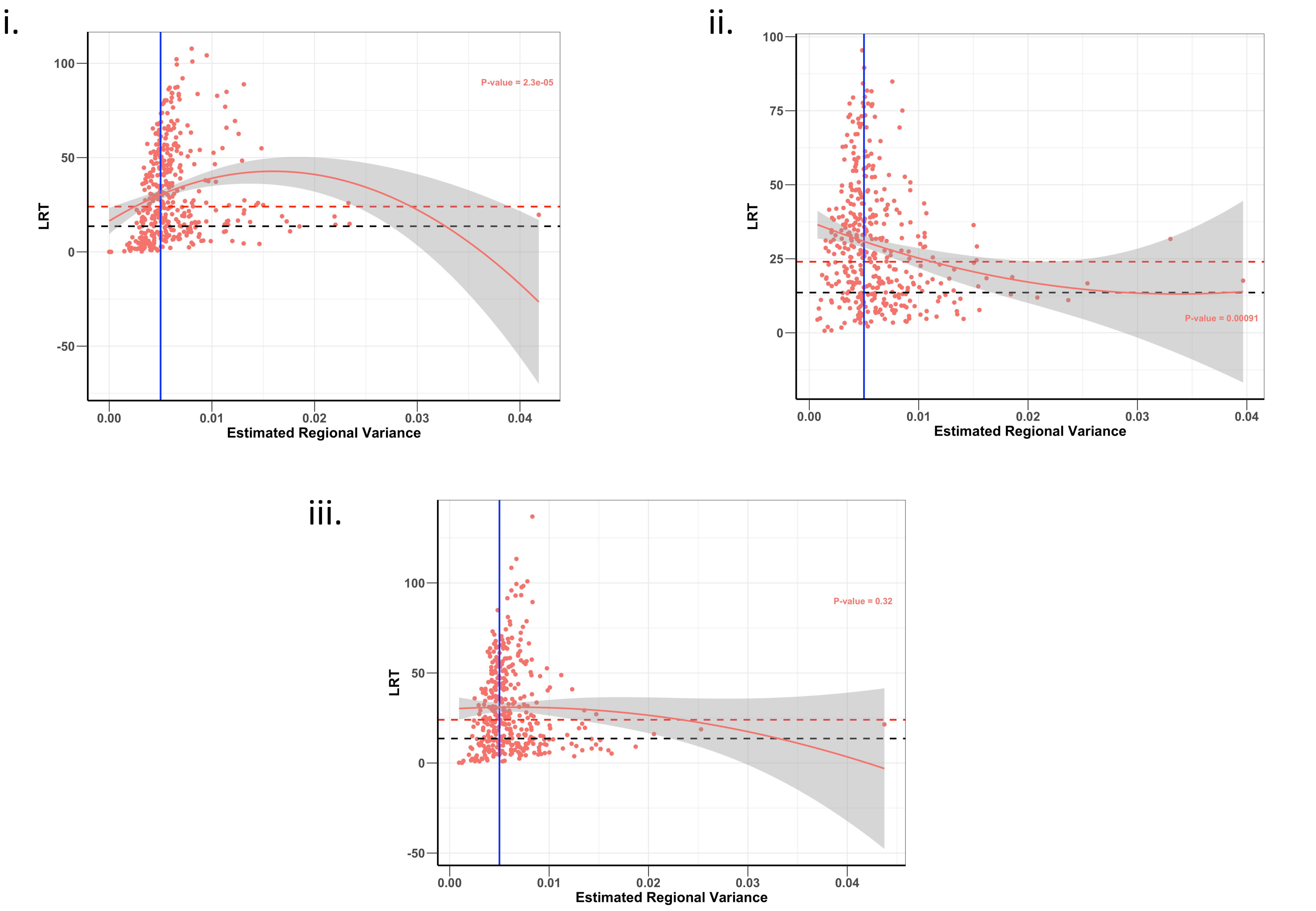


**S6 Fig. Plots of LRT statistic against estimated regional variance for the 20 simulations of each of the three haplotype QTL phenotypes**. The red dashed lines are genome-wide significance threshold (for 48,772 regions), the black dashed lines are Bonferroni significance threshold (for 220 regions), and the blue vertical line is the simulated regional variance of 0.005. The plot (i) is the 1-rare haplotype QTL phenotype, the plot (ii) is the 1-common haplotype QTL phenotype, and the plot (iii) is the multiple haplotype QTL phenotype. The estimated regional variance clusters closely around the simulated value for the regions that pass the Bonferroni threshold.

i.

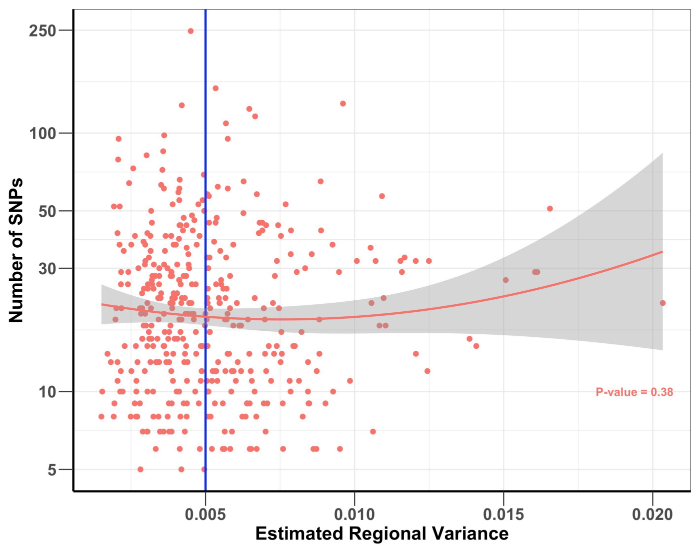


ii.

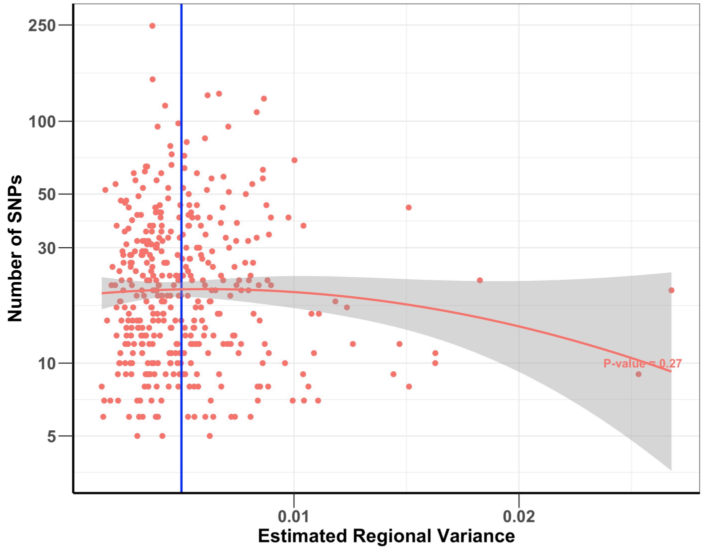


**S7 Fig. Plots of region size against estimated regional variance for the 20 simulations of the two SNP QTL phenotype**. The blue vertical line is the simulated regional variance of 0.005. The plot (i) is the 1-SNP QTL phenotype, and the lower plot (ii) is the multiple SNP QTL phenotype. The two plots show there is no significant relationship between estimated regional variance and region size.

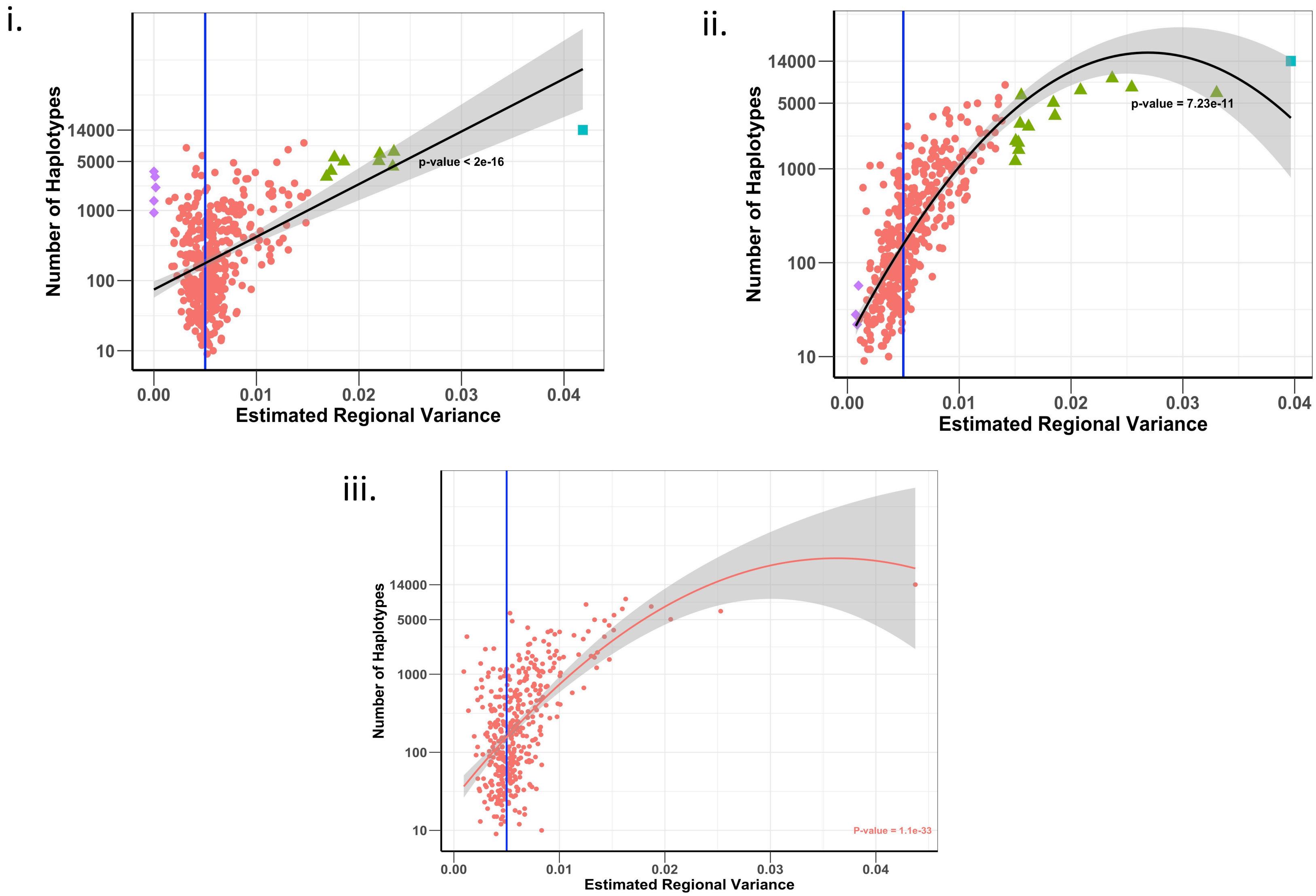


**S8 Fig. Plots of region size against estimated regional variance for the 20 simulations of the three haplotype QTL phenotype**. The blue vertical line is the simulated regional variance of 0.005. The plot (i) is the 1-rare haplotype QTL phenotype, and the plot (ii) is the 1-common haplotype QTL phenotype. On these plots, the blue square point is the region with the largest overestimated variance, the green triangle points are regions with overestimated variance, red points are all other regions and purple points are regions with least variance estimates. The plot (iii) is the multiple haplotype QTL phenotype. The plots show that the estimates of the regional variances are likely to be inflated when the region size gets beyond 1000 haplotypes

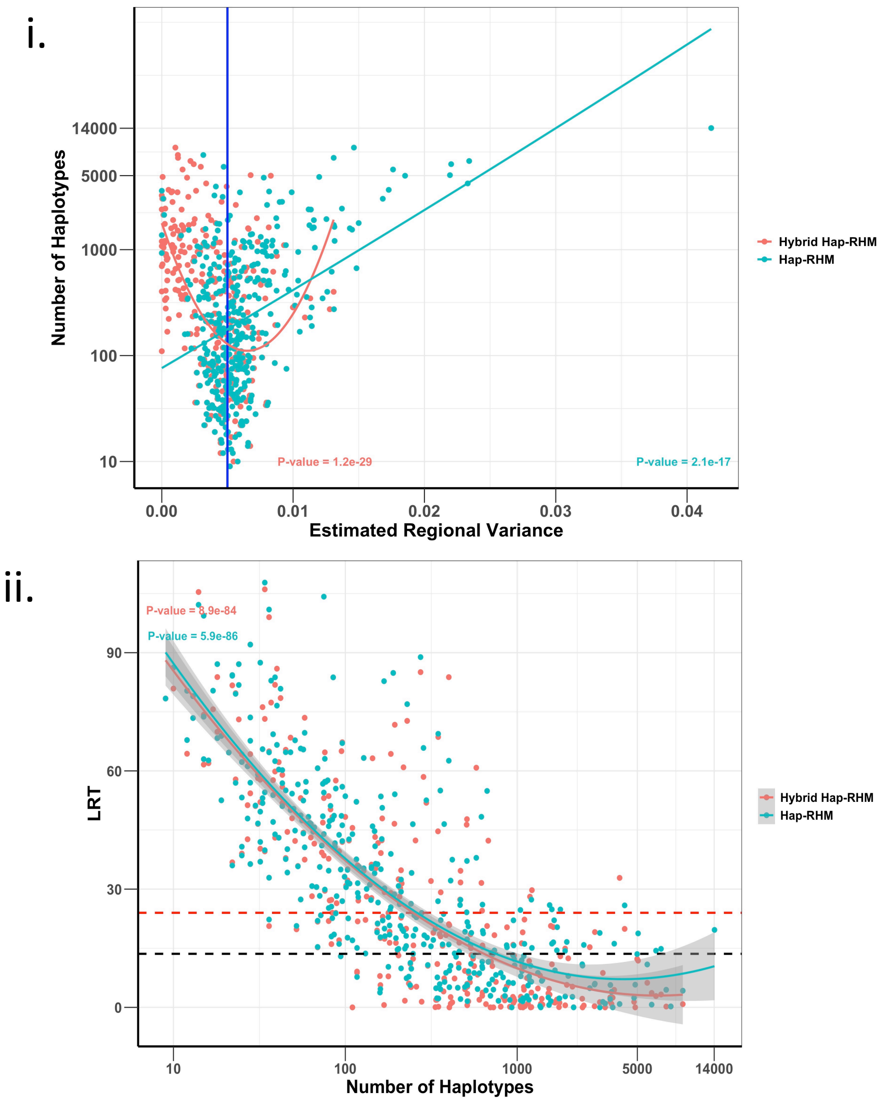


**S9 Fig. Plots for the 1-rare haplotype QTL phenotype analysed using Hap-RHM (red points) and a hybrid variant of the Hap-RHM (blue points).** The hybrid Hap-RHM broke larger regions (regions with more than 20 SNPs) into smaller regions of 20 or less SNPs and used that to determine the haplotypes. Each blue point represents the estimated variances or LRT for best sub-window within the bigger window. (i) is a plot of region size over estimated regional variance for the 20 simulations of the phenotype and (ii) is the plot of LRT statistic over QTL region size for the 20 simulations of each of the phenotype. The plot (i) shows that the hybrid Hap-RHM underestimates the regional variance at larger regions. The LRT statistics however do not improve very much over the default Hap-RHM.

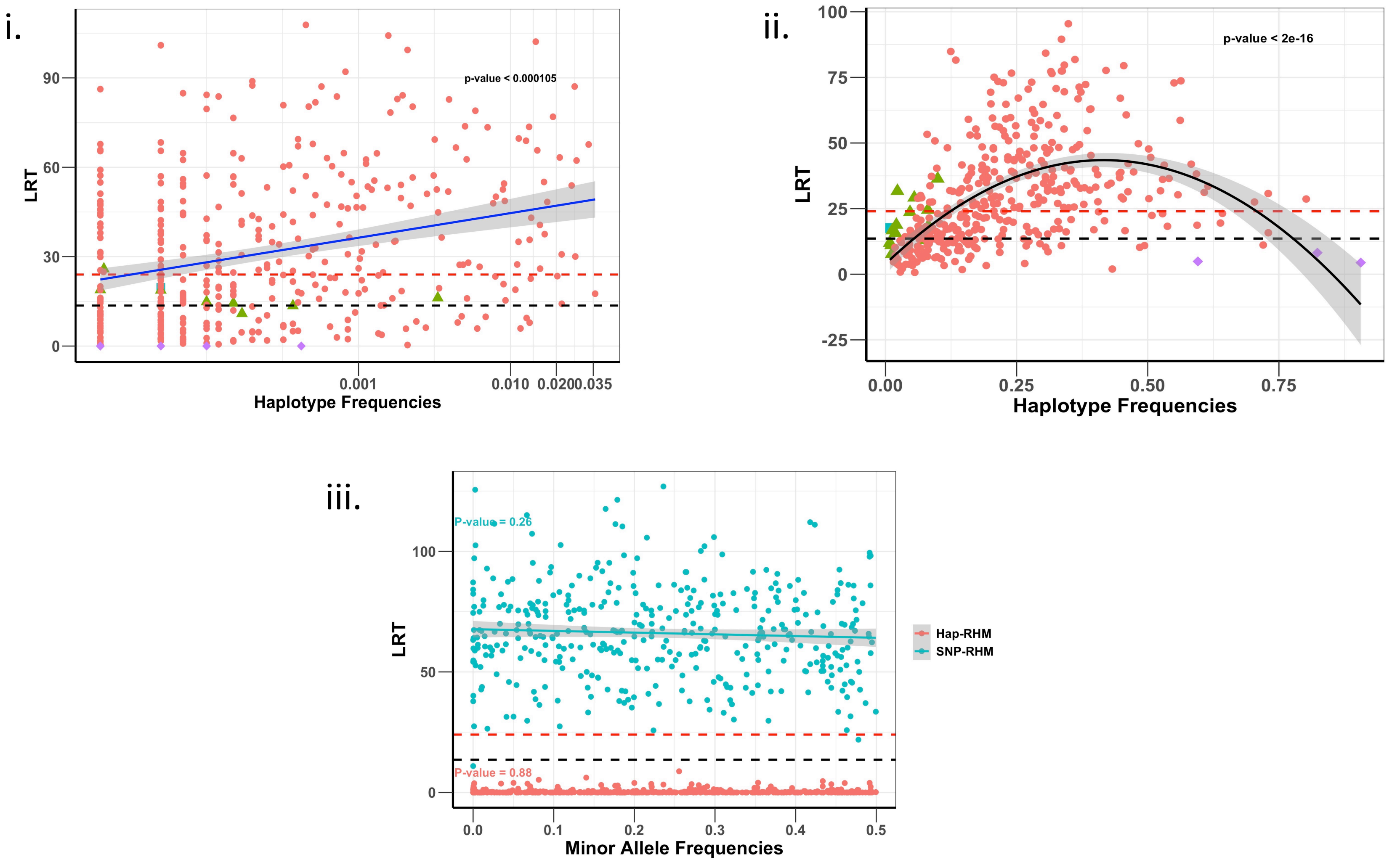


**S10 Fig. Plots of LRT statistic against QTL marker frequencies**. The red dashed lines are genome-wide significance threshold (for 48,772 regions) and the black dashed lines are Bonferroni significance threshold (for 220 regions). The plots (i) and (ii) are 1 rare and 1 common haplotype QTL phenotypes, respectively. On these plots, the blue square point is the region with the largest overestimated variance, the green triangle points are regions with overestimated variance, red points are all other regions and purple points are regions with least variance estimates. The plot (iii) is the 1-SNP QTL phenotype analysed using SNP-RHM (blue points) and Hap-RHM (red points). The three plots show that there is no relationship between QTL marker frequencies and the LRT statistic.

**S1 Table. Top genomic regions identified by SNP/ haplotype-based model for Height.** The columns are chromosome number, the association p-values, the genes within region, genes within 400kb of regions, associated trait, and a list of studies reporting gene association with trait.

| | | | | | **SNP based regional GREML model** | | | | |
|---|---|---|---|---|---|---|---|---|---|
| **CHR** | **P value** | **Genes** | **Genes within 400kb** | **Trait** | **Reference Study** | | | | |
| | | | | | **PMID** | **First author** | **Journal** | **Discovery sample (size and ancestry)** | **Replication sample (size and ancestry)** |
| **1** | 1.78E-05 | | **SPAG17** | height | 18391952 | Weedon M N | Nat Genet | 13,665 European ancestry individuals | 16,482 European ancestry individuals |
| | | | | | 20546612 | Zhao J | BMC Med Genet | 8,184 children of European ancestry | NA |
| **2** | 4.86E-08 | **EFEMP1** | | height | 20881960 | Lango Allen H | Nature | 133,653 European ancestry individuals | 50,074 European ancestry individuals |
| | | | | | 18391952 | Weedon M N | Nat Genet | 13,665 European ancestry individuals | 16,482 European ancestry individuals |
| | | | | | 25282103 | Wood AR | Nat Genet | 253,288 European ancestry individuals | 80,067 European ancestry individuals |
| | | | | | 31217584 | Wojcik GL | Nature | 17,286 African American individuals, 22,192 Hispanic/Latino individuals, 4,680 Asian ancestry individuals, 3,939 Native Hawaiian ancestry individuals, 647 Native American ancestry individuals, | NA |
| | | | **MIR217, MIR216A** | height | 28552196 | Tachmazidou I | Am J Hum Genet | 53,588 European ancestry individuals | 205,003 European ancestry individuals |
| | | | **PNPT1** | height | 25429064 | He M | Hum Mol Genet | 36,227 East Asian ancestry individuals | 57,699 East Asian ancestry individuals |
| | | | | | 20546612 | Zhao J | BMC Med Genet | 8,184 children of European ancestry | NA |
| | | | **SMEK2** | height | 30595370 | Kichaev G | Am J Hum Gene | 458,000 European ancestry individuals | NA |
| **3** | 5.57E-10 | **ZBTB38** | | height | 18391952 | Weedon M N | Nat Genet | 13,665 European ancestry individuals | 16,482 European ancestry individuals |
| | | | | | 25282103 | Wood AR | Nat Genet | 253,288 European ancestry individuals | 80,067 European ancestry individuals |
| | | | | | 23563607 | Berndt SI | Nat Genet | 8,097 European ancestry tall individuals, 8,099 European ancestry short individuals | 4,872 European ancestry tall individuals, 4,831 European ancestry short individuals |
| | | **RASA2** | **RNF7** | height | 28552196 | Tachmazidou I | Am J Hum Genet | 53,588 European ancestry individuals | 205,003 European ancestry individuals |
| | | | **PXYLP1** | height | 28552196 | Tachmazidou I | Am J Hum Genet | 53,588 European ancestry individuals | 205,003 European ancestry individuals |
| | | | **GRK7** | height | 30595370 | Kichaev G | Am J Hum Gene | 458,000 European ancestry individuals | NA |
| **4** | 3.12E-06 | **NCAPG** | | height | 18391951 | Gudbjartsson DF | Nat Genet | 30,968 European ancestry individuals | 8,541 European ancestry individuals |
| | | | | | 20546612 | Zhao J | BMC Med Genet | 8,184 children of European ancestry | NA |
| | | **LCORL** | | height | 22021425 | Carty CL | Hum Mol Genet | 8,149 African American female individuals | Up to 20,809 African American and African ancestry individuals |
| | | | | | 20881960 | Lango Allen H | Nature | 133,653 European ancestry individuals | 50,074 European ancestry individuals |
| | | | | | 18391952 | Weedon M N | Nat Genet | 13,665 European ancestry individuals | 16,482 European ancestry individuals |
| | | | | | 25282103 | Wood AR | Nat Genet | 253,288 European ancestry individuals | 80,067 European ancestry individuals |
| | | | | | 28270201 | Nagy R | Genome Med | 19,920 British ancestry individuals from 6863 families. | NA |

| | | | | | | | | | |
|---|---|---|---|---|---|---|---|---|---|
| | | **DCAF16, FAM184B** | | height | 31217584 | Wojcik GL | Nature | 17,286 African American individuals, 22,192 Hispanic/Latino individuals, 4,680 Asian ancestry individuals, 3,939 Native Hawaiian ancestry individuals, 647 Native American ancestry individuals, | NA |
| | | | **MED28** | height | 30595370 | Kichaev G | Am J Hum Gene | 458,000 European ancestry individuals | NA |
| **4** | 1.98E-06 | **HHIP** | | height | 20881960 | Lango Allen H | Nature | 133,653 European ancestry individuals | 50,074 European ancestry individuals |
| | | | | | 18391952 | Weedon M N | Nat Genet | 13,665 European ancestry individuals | 16,482 European ancestry individuals |
| | | | | | 25282103 | Wood AR | Nat Genet | 253,288 European ancestry individuals | 80,067 European ancestry individuals |
| | | | | | 18391950 | Lettre G | Nat Genet | 15,821 European ancestry individuals | 17,801 European ancestry individuals |
| | | | | | 20546612 | Zhao J | BMC Med Genet | 8,184 children of European ancestry | NA |
| | | | **GYPA** | height | 30595370 | Kichaev G | Am J Hum Gene | 458,000 European ancestry individuals | NA |
| **6** | 3.45E-07 | **GRM4** | | height | 30595370 | Kichaev G | Am J Hum Gene | 458,000 European ancestry individuals | NA |
| | | **HMGA1, MIR6835** | | height | 20881960 | Lango Allen H | Nature | 133,653 European ancestry individuals | 50,074 European ancestry individuals |
| | | | | | 31562340 | Akiyama M | Nat Commun | 159,095 Japanese ancestry individuals | 32,692 Japanese ancestry individuals |
| | | | | | 20546612 | Zhao J | BMC Med Genet | 8,184 children of European ancestry | NA |
| | | **C6orf1, NUDT3** | | height | 19893584 | Kim JJ | J Hum Genet | 8,842 Korean ancestry individuals | NA |
| | | | | | 28270201 | Nagy R | Genome Med | 19,920 British ancestry individuals from 6863 families. | NA |
| | | **RPS10, RPS10–NUDT3** | | height | 31562340 | Akiyama M | Nat Commun | 159,095 Japanese ancestry individuals | 32,692 Japanese ancestry individuals |
| | | | | | 30595370 | Kichaev G | Am J Hum Gene | 458,000 European ancestry individuals | NA |
| | | **PACSIN1** | | height | 30595370 | Kichaev G | Am J Hum Gene | 458,000 European ancestry individuals | NA |
| **6** | 8.53E-09 | **UHRF1BP1, TAF11** | | height | 30595370 | Kichaev G | Am J Hum Gene | 458,000 European ancestry individuals | NA |
| | | **ANKS1A, TCP11, SCUBE3, ZNF76** | | height | 18391951 | Gudbjartsson DF | Nat Genet | 30,968 European ancestry individuals | 8,541 European ancestry individuals |
| | | | | | 20546612 | Zhao J | BMC Med Genet | 8,184 children of European ancestry | NA |
| | | | **C6orf106** | height | 25282103 | Wood AR | Nat Genet | 253,288 European ancestry individuals | 80,067 European ancestry individuals |
| | | | **SPDEF** | height | 28552196 | Tachmazidou I | Am J Hum Genet | 53,588 European ancestry individuals | 205,003 European ancestry individuals |
| | | | **PACSIN1** | height | 30595370 | Kichaev G | Am J Hum Gene | 458,000 European ancestry individuals | NA |
| **6** | 1.21E-05 | **FKBP5** | **RPL10A** | height | 25282103 | Wood AR | Nat Genet | 253,288 European ancestry individuals | 80,067 European ancestry individuals |
| | | **ARMC12, SRPK1, SLC26A8, MAPK14, MAPK13** | | height | 30595370 | Kichaev G | Am J Hum Gene | 458,000 European ancestry individuals | NA |
| | | | **TEAD3, TULP1** | height | 31562340 | Akiyama M | Nat Commun | 159,095 Japanese ancestry individuals | 32,692 Japanese ancestry individuals |

| | | | | | | | | | |
|---|---|---|---|---|---|---|---|---|---|
| | | | **PPARD** | height | 21998595 | N'Diaye A | PLoS Genet | 20,427 African ancestry individuals | 16,436 African American individuals |
| **6** | 9.02E-06 | **BCKDHB** | | height | 25282103 | Wood AR | Nat Genet | 253,288 European ancestry individuals | 80,067 European ancestry individuals |
| **6** | 5.63E-06 | **CENPW** | | height | 25282103 | Wood AR | Nat Genet | 253,288 European ancestry individuals | 80,067 European ancestry individuals |
| | | **RSPO3** | | height | 28270201 | Nagy R | Genome Med | 19,920 British ancestry individuals from 6863 families. | NA |
| **7** | 4.25E-05 | **GATAD1** | | height | 18391951 | Gudbjartsson DF | Nat Genet | 30,968 European ancestry individuals | 8,541 European ancestry individuals |
| | | | | | 20546612 | Zhao J | BMC Med Genet | 8,184 children of European ancestry | NA |
| | | **PEX1** | | height | 25282103 | Wood AR | Nat Genet | 253,288 European ancestry individuals | 80,067 European ancestry individuals |
| | | | | | 20546612 | Zhao J | BMC Med Genet | 8,184 children of European ancestry | NA |
| | | **CDK6, FAM133B** | **KRIT1** | height | 25282103 | Wood AR | Nat Genet | 253,288 European ancestry individuals | 80,067 European ancestry individuals |
| | | | | | 18391952 | Weedon M N | Nat Genet | 13,665 European ancestry individuals | 16,482 European ancestry individuals |
| | | | | | 20546612 | Zhao J | BMC Med Genet | 8,184 children of European ancestry | NA |
| | | | **ANKIB1** | height | 30595370 | Kichaev G | Am J Hum Gene | 458,000 European ancestry individuals | NA |
| **13** | 1.06E-05 | **DLEU1** | **DLEU7, DLEU7–AS1** | height | 28552196 | Tachmazidou I | Am J Hum Genet | 53,588 European ancestry individuals | 205,003 European ancestry individuals |
| | | | | | 25282103 | Wood AR | Nat Genet | 253,288 European ancestry individuals | 80,067 European ancestry individuals |
| | | | | | 18391952 | Weedon M N | Nat Genet | 13,665 European ancestry individuals | 16,482 European ancestry individuals |
| | | | | | 18391951 | Gudbjartsson DF | Nat Genet | 30,968 European ancestry individuals | 8,541 European ancestry individuals |
| | | | | | 20546612 | Zhao J | BMC Med Genet | 8,184 children of European ancestry | NA |
| | | | | | 20881960 | Lango Allen H | Nature | 133,653 European ancestry individuals | 50,074 European ancestry individuals |
| | | | **RNASEH2B–AS1, RNASEH2B** | height | 30595370 | Kichaev G | Am J Hum Gene | 458,000 European ancestry individuals | NA |
| **15** | 7.39E-07 | **ADAMTSL3, SH3GL3** | | height | 18391951 | Gudbjartsson DF | Nat Genet | 30,968 European ancestry individuals | 8,541 European ancestry individuals |
| | | | | | 25282103 | Wood AR | Nat Genet | 253,288 European ancestry individuals | 80,067 European ancestry individuals |
| | | | | | 18391952 | Weedon M N | Nat Genet | 13,665 European ancestry individuals | 16,482 European ancestry individuals |
| | | | | | 20546612 | Zhao J | BMC Med Genet | 8,184 children of European ancestry | NA |
| **15** | 1.15E-05 | **ACAN** | **MRPS11, HAPLN3, ISG20, MRPL46** | height | 20881960 | Lango Allen H | Nature | 133,653 European ancestry individuals | 50,074 European ancestry individuals |
| | | | | | 25282103 | Wood AR | Nat Genet | 253,288 European ancestry individuals | 80,067 European ancestry individuals |
| | | | | | 18391952 | Weedon M N | Nat Genet | 13,665 European ancestry individuals | 16,482 European ancestry individuals |
| | | | | | 21998595 | N'Diaye A | PLoS Genet | 20,427 African ancestry individuals | 16,436 African American individuals |
| | | | | | 20546612 | Zhao J | BMC Med Genet | 8,184 children of European ancestry | NA |
| | | | | | 28552196 | Tachmazidou I | Am J Hum Genet | 53,588 European ancestry individuals | 205,003 European ancestry individuals |
| | | | **FANCI, RLBP1** | height | 25282103 | Wood AR | Nat Genet | 253,288 European ancestry individuals | 80,067 European ancestry individuals |
| | | | **DET1, AEN** | height | 25282103 | Wood AR | Nat Genet | 253,288 European ancestry individuals | 80,067 European ancestry individuals |
| | | | | | 30595370 | Kichaev G | Am J Hum Gene | 458,000 European ancestry individuals | NA |

| | | | | | | | | | |
|---|---|---|---|---|---|---|---|---|---|
| **18** | 2.39E-08 | **CABLES1** | | height | 23563607 | Berndt SI | Nat Genet | 8,097 European ancestry tall individuals, 8,099 European ancestry short individuals | 4,872 European ancestry tall individuals, 4,831 European ancestry short individuals |
| | | | | | 18391951 | Gudbjartsson DF | Nat Genet | 30,968 European ancestry individuals | 8,541 European ancestry individuals |
| | | | | | 20881960 | Lango Allen H | Nature | 133,653 European ancestry individuals | 50,074 European ancestry individuals |
| | | | | | 20546612 | Zhao J | BMC Med Genet | 8,184 children of European ancestry | NA |
| **20** | 1.03E-12 | **UQCC1, GDF5** | **FAM83C−AS1, FAM83C, CPNE1, RBM12, NFS1, RBM39, ROMO1, MYH7B, TRPC4AP** | height | 18391951 | Gudbjartsson DF | Nat Genet | 30,968 European ancestry individuals | 8,541 European ancestry individuals |
| | | | | | 18391952 | Weedon M N | Nat Genet | 13,665 European ancestry individuals | 16,482 European ancestry individuals |
| | | | | | 28270201 | Nagy R | Genome Med | 19,920 British ancestry individuals from 6863 families. | NA |
| | | | | | 18193045 | Sanna S | Nat Genet | 6669 European ancestry individuals | 3860 African American or Afro-Caribbean 23684 European ancestry individuals |
| | | | | | 31562340 | Akiyama M | Nat Commun | 159,095 Japanese ancestry individuals | 32,692 Japanese ancestry individuals |
| | | | | | 30595370 | Kichaev G | Am J Hum Gene | 458,000 European ancestry individuals | NA |
| | | | | | 25429064 | He M | Hum Mol Genet | 36,227 East Asian ancestry individuals | 57,699 East Asian ancestry individuals |
| | | **CEP250** | | height | 18391951 | Gudbjartsson DF | Nat Genet | 30,968 European ancestry individuals | 8,541 European ancestry individuals |
| | | | **MMP24, EIF6** | height | 18391951 | Gudbjartsson DF | Nat Genet | 30,968 European ancestry individuals | 8,541 European ancestry individuals |
| | | | **ERGIC3, FER1L4** | height | 28552196 | Tachmazidou I | Am J Hum Genet | 53,588 European ancestry individuals | 205,003 European ancestry individuals |
| | | | **PHF20** | height | 25282103 | Wood AR | Nat Genet | 253,288 European ancestry individuals | 80,067 European ancestry individuals |
| | | | **PROCR** | height | 23563607 | Berndt SI | Nat Genet | 8,097 European ancestry tall individuals, 8,099 European ancestry short individuals | 4,872 European ancestry tall individuals, 4,831 European ancestry short individuals |
| | | | **EDEM2** | height | 25282103 | Wood AR | Nat Genet | 253,288 European ancestry individuals | 80,067 European ancestry individuals |
| **20** | 7.91E-08 | **CEP250, ERGIC3, FER1L4, CPNE1, RBM12, NFS1, ROMO1, RBM39** | **GDF5** | height | 18391951 | Gudbjartsson DF | Nat Genet | 30,968 European ancestry individuals | 8,541 European ancestry individuals |
| | | | | | 28552196 | Tachmazidou I | Am J Hum Genet | 53,588 European ancestry individuals | 205,003 European ancestry individuals |
| | | | | | 31562340 | Akiyama M | Nat Commun | 159,095 Japanese ancestry individuals | 32,692 Japanese ancestry individuals |
| | | | | | 18391952 | Weedon M N | Nat Genet | 13,665 European ancestry individuals | 16,482 European ancestry individuals |
| | | **PHF20, CNBD2** | | height | 25282103 | Wood AR | Nat Genet | 253,288 European ancestry individuals | 80,067 European ancestry individuals |
| | | **EPB41L1** | | height | 23563607 | Berndt SI | Nat Genet | 8,097 European ancestry tall individuals, 8,099 European ancestry short individuals | 4,872 European ancestry tall individuals, 4,831 European ancestry short individuals |

#### Haplotype bases regional GREML model

| | | | | | | | | | |
|---|---|---|---|---|---|---|---|---|---|
| **6** | 1.96E-05 | **SAMD5** | | height | 30595370 | Kichaev G | Am J Hum Gene | 458,000 European ancestry individuals | NA |

**S2 Table. Top genomic regions identified by SNP/ haplotype-based model for MDD**. The columns are chromosome number, the association p-values, the genes within region, genes within 400kb of regions, associated trait, and a list of studies reporting gene association with trait.

| CHR | P value | Genes | Genes within 400kb | Traits | PMID | First author | Journal | Discovery sample (size and ancestry) | Replication sample (size and ancestry) |
|---|---|---|---|---|---|---|---|---|---|
| **SNP based regional GREML model** | | | | | | | | | |
| **13** | 2.41E-05 | | **DACH1** | depressive symptom measurement, stressful life event measurement | 30718454 | Arnau-Soler A | Transl Psychiatry | 4919 European ancestry individuals | NA |
| | | | | age at onset of Alzheimer's disease | 26830138 | Herold C | Mol Psychiatry | 3524 European ancestry individuals | NA |
| | | | | brain volume measurement | 31676860 | Zhao B | Nat Genet | 19629 European ancestry individuals | NA |
| | | | | schizophrenia, attempted suicide | 31164008 | Mullins N | Am J Psychiatry | 8764 European ancestry individuals | NA |
| **Haplotype bases regional GREML model** | | | | | | | | | |
| **18** | 2.30E-06 | | **DCC** | unipolar depression | 29700475 | Wray NR | Nat Genet | 480359 European ancestry individuals | NA |
| | | | | | 30718901 | Howard DM | Nat Neurosci | 446238 European ancestry individuals | 1306354 European ancestry individuals |
| | | | | | 27422368 | Zeng Y | Biol Psychiatry | 21,387 European ancestry individuals | 18,759 European ancestry individuals |
| | | | | self-reported educational attainment, mathematical ability, cognitive function measurement | 29186694 | Lam M | Cell Rep | 436124 European ancestry individuals | NA |
| | | | | | 33414549 | Demange PA | Nat Genet | 257700 European ancestry individuals | NA |
| | | | | | 27225129 | Okbay A | Nature | 405072 European ancestry individuals | NA |
| | | | | | 30038396 | Lee JJ | Nat Genet | 1131438 European ancestry individuals | NA |
| | | | | Neuroticism measurement | 29292387 | Turley P | Nat Genet | 168105 European ancestry individuals | NA |
| | | | | | 30643256 | Baselmans BML | Nat Genet | 523783 European ancestry individuals | 59206 European ancestry individuals |
| | | | | brain volume measurement | | Hibar DP | Nature | 13171 European ancestry individuals | 736 Hispanic or Latin American<br>15130 European<br>545 East Asian |

| | | | | | | | | | |
|---|---|---|---|---|---|---|---|---|---|
| **3** | 5.19E-06 | **MYRIP** | **EIF1B–AS1** | Alzheimer's disease | 20197096 | Stein JL | Neuroimage | 742 European ancestry individuals | NA |
| | | | | sleep measurement | 17903308 | Gottlieb DJ | BMC Med Genet | 738 NR | NA |
| | | | | reading and spelling ability | 30741946 | Gialluisi A | Transl Psychiatry | 2,563 European ancestry individuals | NA |
| | | | | information processing speed | 21130836 | Luciano M | Biol Psychol | 4,039 European ancestry individuals | NA |
| | | | | Grey matter density measurement | 31530798 | Alliey-Rodriguez N | Transl Psychiatry | 483 European ancestry, 44 NR, 250 African American or Afro-Caribbean ancestry | NA |
| | | | | white matter hyperintensity measurement | 32517579 | Armstrong NJ | Stroke | 16,832 European ancestry individuals, 736 African American individuals, 658 Hispanic individuals | 8,428 European and unknown ancestry individuals |
| | | | **MOBP** | Alzheimer's disease (cognitive decline) | 23535033 | Sherva R | Alzheimers Dement | 303 European ancestry cases | NA |
| | | | | Cognitive performance | 19734545 | Need AC | Hum Mol Genet | 958 European ancestry, 207 East Asian ancestry, 26 NR, 104 South Asian ancestry | NA |
| | | | | Progressive supranuclear palsy | 21685912 | Höglinger GU | Nat Genet | 4027 European ancestry, 45 NR, 329 Other | 4611 European ancestry individuals |
| | | | | Attention deficit hyperactivity disorder | 18839057 | Lesch KP | J Neural Transm | 593 European ancestry individuals | NA |
| **2** | 2.07E-05 | **ANO7** | **PASK, MTERFD2, SNED1** | Unipolar depression, schizophrenia, bipolar disorder, concentration dose ratio, response to antipsychotic drug | 25944848 | Athanasiu L | J Psychopharmacol | 829 European ancestry cases | NA |
| | | | **BOK, THAP4** | brain measurement | 20171287 | Stein JL | Neuroimage | 740 European ancestry individuals | NA |
| | | | **PPP1R7** | PHF-tau measurement | 32450446 | Wang H | Neurobiol Aging | 1285 NR | NA |
| **1** | 2.46E-05 | **KCNK1** | **KIAA1804** | age at onset Alzheimer's disease, | 26830138 | Herold C | Mol Psychiatry | 3524 European ancestry individuals | NA |
| | | | | cognitive function measurement | 28808816 | Xu C | Biogerontology | 278 East Asian ancestry individuals | NA |
| **12** | 2.70E-05 | **AMIGO2, PCED1B** | | Alzheimer's disease | 19118814 | Beecham GW | Am J Hum Genet | 988 European ancestry individuals | 458 European ancestry individuals |
| | | | | PHF-tau measurement | 32450446 | Wang H | Neurobiol Aging | 1285 NR | NA |

| | | | | | | | | | |
|---|---|---|---|---|---|---|---|---|---|
| | | | | bipolar disorder, attempted suicide | 31164008 | Mullins N | Am J Psychiatry | 8764 European ancestry individuals | NA |
| **10** | 2.82E-05 | | **NRP1** | unipolar depression, alcohol dependence | 29071344 | Zhou H | JAMA Psychiatry | 3,041 African American individuals, 1,618 European ancestry individuals | 1,612 African American individuals, 1,551 European ancestry individuals |
| | | | | schizophrenia, response to paliperidone, schizophrenia symptom severity measurement | 27846195 | Li Q | Pharmacogenet Genomics | 1,390 European ancestry cases | NA |
| | | | | migraine disorder | 23793025 | Anttila V | Nat Genet | 118710 European ancestry individuals | NA |
| | | | | brain measurement | 32665545 | van der Meer D | Nat Commun | 26,502 European ancestry individuals | NA |
| | | | **LINC00838** | Schizophrenia | 21682944 | Alkelai A | Int J Neuropsychopharmacol | 331 European ancestry individuals | 189 Greater Middle Eastern ancestry individuals |
| **2** | 4.59E-05 | | **HNRNPA3, MIR4444−2, MIR4444−1, NFE2L2, MIR3128, LOC100130691, MIR6512** | alcohol consumption measurement | 30643251 | Liu M | Nat Genet | 1,039,210 European ancestry individuals | NA |
| **2** | 4.67E-05 | **CRIM1** | **FEZ2** | cognitive function measurement, mathematical ability | 30038396 | Lee JJ | Nat Genet | 1131438 European ancestry individuals | NA |
| | | | | intelligence | 29844566 | Davies G | Nat Commun | 300,486 European ancestry individuals | NA |
| | | | **VIT** | brain volume measurement | 31676860 | Zhao B | Nat Genet | 19629 European ancestry individuals | NA |
